# Proteasome inhibition reprograms chromatin landscape in breast cancer

**DOI:** 10.1101/2023.10.13.562284

**Authors:** H Karimi Kinyamu, Brian D. Bennett, James M. Ward, Trevor Archer

## Abstract

The 26S proteasome is the major protein degradation machinery in cells. Cancer cells use the proteasome to modulate gene expression networks that promote tumor growth. Proteasome inhibitors have emerged as effective cancer therapeutics, but how they work mechanistically remains unclear. Here, using integrative genomic analysis, we discovered unexpected reprogramming of the chromatin landscape and RNAPII transcription initiation in breast cancer cells treated with the proteasome inhibitor MG132. The cells acquired dynamic changes in chromatin accessibility at specific genomic loci termed Differentially Open Chromatin Regions (DOCRs). DOCRs with decreased accessibility were promoter proximal and exhibited unique chromatin architecture associated with divergent RNAPII transcription. Conversely, DOCRs with increased accessibility were primarily distal to transcription start sites and enriched in oncogenic super enhancers predominantly accessible in non-basal breast tumor subtypes. These findings describe the mechanisms by which the proteasome modulates the expression of gene networks intrinsic to breast cancer biology.

**Highlights:**

- Proteasome inhibition uncovers *de novo* Differential Open Chromatin Regions (DOCRs) in breast cancer cells.
- Proteasome inhibitor sensitive promoters exhibit a distinctive chromatin architecture with discrete transcription initiation patterns.
- Proteasome inhibition reprograms accessibility of super enhancers.
- Proteasome inhibitor sensitive super enhancers distinguish basal from non-basal breast cancer subtypes.

## INTRODUCTION

The 26S proteasome is a large multi-enzymatic complex, which regulates cellular protein homeostasis by selective degradation of ubiquitinated proteins (1). Proteasome activity is required for many cellular functions and in particular, the proteasome directly regulates chromatin structure and function to influence transcription and gene expression.

Regulation of chromatin function is critical to epigenetic mechanisms that control transcription and gene expression. Chromatin is used by cells to package DNA in the nucleus, a complex process achieved by the compaction of DNA with histone proteins to form nucleosomes, which are the basic units of chromatin (2,3). The packaging of DNA into chromatin is a key regulatory step for controlling DNA templated processes including transcription (4). Chromatin function is in part dictated by the underlying DNA sequence, and in this regard several processes such as nucleosome remodeling through post-translational modifications of histones and eviction of nucleosomes to expose underlying regulatory DNA elements, can alter the physical properties of chromatin that typically either enhance or repress transcription (4-6).

The proteasome regulates chromatin function at many levels. The complexity of this regulation involves both proteolytic and non-proteolytic activities of the proteasome (7). The expression of many proteins that regulate chromatin dynamics such as chromatin remodeling complexes and histone modifying enzymes are tightly regulated by proteasomal degradation (8). Further recent evidence suggesting histone proteins are direct targets of proteasomal degradation, underscores a critical role for the proteasome in fine tuning chromatin architecture and function (9).

The intimate connection between the proteasome and transcriptional regulation is dynamic and several studies suggest a multifaceted function of the proteasome in transcription regulation that may involve proteolytic and non-proteolytic activities of proteasome complex (10). Non-proteolytic proteasome activities in transcription are supported by studies showing proteasome complex subunits are nuclear proteins and components of the RNA polymerase II (RNAPII) complexes (11-16). The discovery that transcription factor activation domains often overlap degradation signals led support for a direct role for proteolytic activities of the proteasome in transcriptional regulation (17). Furthermore, RNAPII itself is a major target of proteasomal degradation, thereby impacting the transcriptional process at every step (18).

Altered chromatin function is linked to aberrant transcriptional regulation in cancer (19,20). Cancer cells often possess elevated levels of proteasome activity and are therefore more sensitive than normal cells to proteasome inhibitors (21). Proteasome inhibitor drugs have been used for several years to treat several hematological malignancies, and clinical studies testing their effectiveness in solid tumor cancers are ongoing (22,23). Their mode of drug action is generally perceived to be via protein turnover, however other mechanisms may be at play. A fuller understanding of gene networks that converge to control tumor cell growth and survival in cells exposed to proteasome inhibitors is critical to harnessing their therapeutic potential.

In a recent study, we showed treatment of MCF7 breast cancer cells with proteasome inhibitor MG132 induced and repressed the spreading of H3 lysine methylation and acetylation at promoters of genes that encode tumor suppressors and cell proliferation pathways, respectively (24). Here, we assessed chromatin accessibility, epigenome, and transcriptome dynamics to uncover regions of the breast cancer cell genome that were sensitive to proteasome inhibition. We have uncovered profound changes in chromatin accessibility, epigenome, and transcriptome dynamics in breast cancer cells in response to proteasome inhibition. We found that proteasome activity was required for the accessibility of specific cis-regulatory elements in the MCF-7 breast cancer cell genome. Changes in accessible chromatin in treated cells were associated with enriched active histone marks and differences in RNAPII transcription. Chromatin changes distal to gene promoters upon MG132 treatment underscore the proteasome’s role in regulating chromatin state and RNAPII transcription from cis-regulatory elements critical for tumor maintenance.

## MATERIALS AND METHODs

### Experimental model system and treatment conditions

#### Cell line and cell culture

MCF-7 (ATCC® HTB-22™) breast cancer cells were routinely maintained in a humidified incubator at 37°C with 5% CO2 in Modified Eagle Medium (MEM, GIBCO) supplemented with 10% fetal bovine serum (FBS, Atlanta Biologicals), 100 μg/mL penicillin/streptomycin, 2 mM glutamine and 10 mM HEPES (GIBCO).

#### Experimental conditions for inhibiting proteasome activity

MG132 is a small molecule that effectively blocks the proteolytic activity of the 26S proteasome complex. To inhibit proteasome activity, MCF-7 breast cancer cells were seeded for 24 hours in phenol red-free MEM medium (GIBCO) supplemented with 5% charcoal-stripped calf serum (Atlanta Biologicals), 10 mM HEPES and 2 mM glutamate. After 24 hours, cells were switched to medium containing vehicle (DMSO, Sigma) as control or 1 uM MG132 (Calbiochem) for 4 and 24 hours.

### Transcription and gene expression analysis

#### RNA extraction

Total RNA was extracted from three biological replicates of vehicle and MG132 treated cells using total RNA isolation kit (Norgen Biotek, Thorold, ON, Canada). RNA concentration was determined using Nanodrop ND-1000 spectrophotometer (Thermo Fisher Scientific). The integrity of the RNA sample was verified using the Agilent RNA 6000 kit and Agilent Bioanalyzer 2100 (Agilent Technologies). Samples with an RNA Integrity Number (RIN) equal or above 8 were considered appropriate for further downstream analysis.

#### mRNA expression quantification by RNA-Sequencing (RNA-Seq)

RNA-seq of ribosomal depleted RNA was performed by Expression Analysis, Q^2^ Solutions (Morrisville, NC). Libraries were sequenced to a depth of an average of 60 million reads (paired-end, 2×75 bp). Raw reads were quality filtered to include only those with a mean Phred quality score of 20 or greater. Adapter was trimmed using Cutadapt version 1.12. The preprocessed reads were aligned to the hg19 assembly using STAR version 2.6.0 (25). The STAR index was built using a GTF file derived from GENOCDE v27 and --sjdbOverhang 10. Read counts were generated using the feature Counts tools from the Subread package version 1.5.1. Differentially expressed genes (log2 fold change of ±1; adjusted p-value ≤ 0.05) were detected with the DESeq2 package (26).

#### Analysis of RNA Pol II genome occupancy to quantify transcriptional activity

RNA polymerase II (RNA pol II) genome occupancy was used as a surrogate to measure transcription rates across the genome. Chromatin immunoprecipitation and sequencing assays (ChIP-seq) were performed to identify regions of the genome enriched with RNA Pol II in MCF-7 cells treated with MG132. Cells were treated with vehicle or MG132 as specified above, and chromatin immunoprecipitation (ChIP) for RNA Pol II done as follows: MCF-7 cells were briefly cross-linked in 1% formaldehyde phosphate buffered saline (PBS) for 5 minutes. After 5 minutes the cross link was quenched with glycine (125 mM) for 5 minutes. Cells were pelleted by centrifugation and resuspended in sonication buffer (20 mM Tris 8.0, 150 mM NaCl, 0.5% Triton X-100, 2 mM EDTA, 10% glycerol) supplemented with protease inhibitors and incubated on ice for 10 minutes. Chromatin was fragmented in 15 mL tubes using the bioruptor (Diagenode) for 12 cycles (30 sec on/30 sec off), total sonication time 6 minutes. After sonication the fragmented chromatin was spun briefly in a cooled centrifuge, transferred to 1.5 mL Eppendorf tubes, and centrifuged at 14000 RPM for 10 minutes. Following centrifugation an aliquot of the chromatin was diluted with immunoprecipitation buffer and incubated with various RNA Pol II antibodies for 12-14 hours at 4°C. DNA/protein immunocomplexes were recovered as described below for histone modification ChIP.

### Analysis of annotated and nonannotated transcription from cis-regulatory elements

#### Start-seq analysis to detect short capped RNAs

Start-seq quantifies short, capped, chromatin-associated RNAs, which are indicative of newly synthesized nascent RNA. It is a useful method for detecting transcription from cis-regulatory elements, including promoter and enhancer elements. Short capped RNAs were prepared as previously described (27) with some modifications (28). Approximately 20 million nuclei were isolated from control and MG132 treated cells using hypotonic lysis buffer followed by total RNA extraction using Trizol reagent (Invitrogen). Small RNAs, 25-80 nt, were size selected on 15% Urea-TBE gel (Novex). Gel slices were crushed by centrifugation (16,000 x g) and RNA eluted following incubation in 300 mM NaCl at room temperature for 2.5 hours. The eluted sized RNA was separated from the gel slices by short centrifugation using cellulose acetate spin filters (Agilent cat# 5185-5990). The 5’-triphosphates on these RNAs were converted to monophosphates by treating the short nuclear RNA 5’ Polyphosphatase. The digestion reaction was purified using Zymo Oligo clean and concentrator kit. The eluted RNA was then treated with 5’ Terminator exonuclease enzyme (Epicentre) to remove the uncapped RNA, and capped RNAs were recovered using the ZYMO kit as above. Sequencing libraries were prepared using Illumina TruSeq Small RNA as per the manufacturer’s instructions, with the ligation of 3′-small RNA Tru-Seq adapter using the truncated T4 RNA Ligase 2 (NEB). 45-150 nt capped small RNAs were recovered on 15% Urea-TBE gel (Novex) and RNA was extracted from the gel as described above. The RNA 5’ ends were dephosphorylated using Heat Labile Alkaline phosphatase (Epicentre), purified using ZYMO columns and treated with RNA 5’ Pyrophosphohydrolase (RppH) to create 5’ monophosphate RNA, followed by another column clean up and elution of 5’ ends dephosphorylated RNA. For small RNA sequencing, 5′-Tru-Seq small RNA adapters were ligated on to the de-capped RNA with T4 RNA Ligase 1 (NEB) in the presence of ATP and cDNA synthesized following Illumina small RNA-seq protocol. Small RNAs we enriched by PCR amplification and size separated on 6% native gel (Novex). Gel slices corresponding to RNA sizes between 100 and 200 bp were excised, and RNA purified using Qiagen MinElute kit. Concentration of the library was determined using Qubit and RNA sequenced using NextSeq 500 system (Illumina)..

### Analysis of chromatin state

#### Chromatin accessibility

Assay for transposase-accessible chromatin and sequencing (ATAC-seq) ATAC-seq uses a hyperactive Tn5 transposase to insert Illumina sequencing adaptors into accessible chromatin regions in the genome. ATAC-seq is a powerful approach to identify potential regulatory elements in the whole genome. To perform ATAC-seq, MCF-7 nuclei were prepared as originally described with minor modifications (29). Briefly, 50,000 cells were pelleted and resuspended in ice-cold cytoskeletal (CSK) lysis buffer, 10 mM PIPES pH 6.8, 100 mM NaCl, 300 mM sucrose, 3 mM MgCl2, 0.1% TritonX-100 (30). Cells were incubated on ice for 5 minutes and spun at 500Xg for 5 minutes at 4 °C, followed by digestion with 5 uL of transposase (Nextra DNA kit) (29,30). Transposase digestion was performed for 30 minutes at 37 °C, on a block shaker set at 200 rpm. Digested DNA was purified using MinElute PCR purification kit (Qiagen, Frederick, MD) and eluted from the MinElute columns with a 10 μL volume of Elution Buffer (Qiagen). Accessible DNA libraries were amplified by PCR using NEBNext High-Fidelity PCR Master Mix (New England Biolabs, Ipswich, MA), custom Nextera PCR primers as previously described (29). Libraries were amplified for a total of 10 PCR cycles after verifying in each case the number of cycles for optimal amplification. DNA purification and size selection were performed using AmpureXP SPRI select beads (Beckman Coulter, Indianapolis, IN). Fragmented DNA was eluted in 30 µl elution buffer (Qiagen PCR kit). An aliquot of the DNA was resolved on 1% agrose gel and visualized on ChemiDoc Touch Imaging System (Biorad) to determine the quality of the tagmented libraries. Samples were subjected to paired-end sequencing using 2 × 50 bp reads on Illumina NextSeq High2000 at the NIEHS Epigenomics core (Research Triangle Park, NC).

### Analysis of epigenetic modifications

#### Detection of histone modification patterns

Chromatin immunoprecipitation and sequencing assays (ChIP-seq) were performed to identify regions of the genome enriched with specific histone modifications in MCF-7 cells treated with MG132. Cells were treated with vehicle or MG132 as specified above. Chromatin immunoprecipitation for histone modifications was done as follows: MCF-7 cells were briefly cross-linked in 1% formaldehyde phosphate buffered saline (PBS) for 5 minutes. After 5 minutes the cross link was quenched with glycine (125 mM) for 5 minutes. The quenched cross linker was quickly discarded, and cells rinsed twice with PBS supplemented with protease inhibitors. Cells were scraped and harvested in 5 mL PBS with an additional 5 mL PBS rinse to collect majority of the cells. The cells were pelleted by centrifugation at 4°C, followed by nuclei isolation as previously described (31). To fragment chromatin, nuclei were resuspended in micrococcal nuclease (MNase, Worthington Enzymes) digestion buffer (10 mM Tris-HCl pH 7.4, 15 mM NaCl, 60 mM KCl, 1 mM Cacl2, 0.15mM spermidine, 0.5mM spermidine) on ice. Nuclei were digested with 50 units of MNase nuclease on a temperature-controlled heating block at 25°C for 20 minutes with gentle mixing at 400 rpm. Following digestion, samples were quickly placed on ice and the reaction stopped by addition of EDTA/EGTA stop buffer (100 mM, EDTA, 10 mM EGTA, pH 7.5), gently mixed by pipetting, followed by addition of SDS lysis buffer (4%SDS, 40 mM EDTA, 200 mM Tris pH 8.0), final concentration 1% SDS supplemented with Halt™ Protease Inhibitor Cocktail (Thermo Fisher). To further release chromatin, nuclei were briefly disrupted using a mini homogenizer for 5 sec and further incubated on the Hulamixer sample mixer (Invitrogen) for 10 minutes at 4°C. Chromatin was recovered by centrifugation at 14,000 rpm for 10 minutes. Following centrifugation an aliquot of MNase fragmented chromatin was diluted 10X with immune precipitation buffer (20 mM Tris 8.0, 150 mM NaCl, 0.5% Triton X-100, 2 mM EDTA, 10% glycerol) supplemented with protease inhibitors (Roche). Antibodies against histone modifications of interest were added and chromatin incubated overnight at 4°C on a slow rotating nutator. The next day, 20 µl Protein A and G magnetic Dynabeads (Invitrogen) were added, and samples were incubated for an additional 2 hours at 4°C on nutator to capture DNA/protein immunocomplexes. Following incubation, protein/ DNA immunocomplexes were recovered by subsequent washes and eluted as previously described (32). Eluted immunoprecipitated complexes were digested with RNAse A (Qiagen) followed by proteinase K digestion and reverse cross linking as previously described (32). DNA was recovered using Qiagen PCR kit purification system (Qiagen) and quantified using Quant-iT^TM^ dsDNA HS assay kit with Qubit^TM^ fluorometer (Invitrogen).

### ChIP sequencing library preparation

Following DNA recovery from RNA Pol II and histone modification immunocomplexes, libraries were prepared with Ilumina compatible NEXTflex Rapid DNA-seq Kit and sequenced on a NextSeq2000 (Illumina). At least two independent biological replicates were performed for each histone modification and RNA Pol II chromatin immunoprecipitation assays.

### ChIP-Seq processing

The FASTQ files for each biological replicate were concatenated for each sample. Raw reads were quality filtered to include only those with a mean Phred quality score of 20 or greater. Adapter was trimmed using Cutadapt version 1.12. The preprocessed reads were aligned to the hg19 assembly using Bowtie version 1.2 and parameters -v 2 -m 1 - -best --strata (33). Aligned reads were deduplicated by only keeping one read pair when multiple pairs had both mates aligned to the same position. Bound locations were obtained from the aligned reads by extracting the entire length of the aligned fragment. Coverage tracks were generated from these bound locations using the genomecov tool from the bedtools suite version 2.17.0 (34). The coverage tracks were normalized to depth per 10 million mapped reads.

### Analysis of Super-Enhancers

Additional peaks were called for the H3K27ac ChIP-seq samples using MACS2 and parameters -q 0.000001 --fe-cutoff 5. Super-enhancers were identified from these peaks using ROSE and parameter -t 2500 (35).

### ATAC-Seq data processing

Raw reads were quality filtered to only include those with a mean Phred quality score of 20 or greater. Adapter was trimmed using Cutadapt version 1.12. The preprocessed reads were aligned to the hg19 assembly using Bowtie version 1.2 and parameters -v 2 -m 1 --best --strata. Reads aligning to the mitochondrial chromosome were removed. Surviving reads were deduplicated by only keeping one read pair when multiple pairs had both mates aligned to the same position. Nucleosome-free reads were extracted by requiring the fragment length to be less than 100 bases. All heatmaps, metaplots, and downstream analyses were done using the coverage from these nucleosome-free reads. Coverage tracks were generated using the genomecov tool from the bedtools suite version 2.17.0. The coverage tracks were normalized to depth per 10 million mapped reads. Peaks were called independently for each sample using MACS2 v2.1.0 and parameters -q 0.0001 --nomodel --extsize 9. Peaks for all samples were merged, and any resulting merged peaks that were within 100 bases of each other were also merged, resulting in a set of global open chromatin regions. Counts for each sample and open chromatin region were obtained by counting the number of reads that overlap with the open chromatin regions. Differential open chromatin regions (DOCRs) were identified using DESeq2, requiring an FDR of less than 0.05 and at least 100 reads in each sample for that open chromatin region. A DOCR was categorized as promoter if its center fell with the region 1 kb upstream to 500 bases downstream of an annotated TSS. Remaining DOCRs were categorized as genic if the center fell within the gene body of an annotated TSS, and all other DOCRs were categorized as intergenic. Epilogos chromatin state scores in 200-base bins were used to define consensus chromatin states with highest score in each bin (36). DOCRs were annotated using the 15 Epilogos chromatin states, based upon overlap with the center of each DOCR. DOCRs were summarized by direction of change, genome category, and chromatin state.

### Heatmap and metaplot processing

Signal used in heatmaps and metaplots (for all data types) was derived from the coverage of aligned fragments. All signal (except for start RNA) was normalized to “depth per 10 million mapped fragments”, by multiplying the coverage signal by 10 million and dividing by the number of aligned fragments. Heatmaps were generated by specifying 100 bins covering the genomic regions in the heatmap, and by calculating the average normalized signal in those genomic regions for each bin. Histone modification ChIP-seq heatmaps were further normalized by first generating a similar heatmap with Total H3 signal, then subtracting the Total H3 signal from the histone modification ChIP-seq signal for each bin. RNAPII ChIP-seq heatmaps were normalized in a similar way, except that input signal was subtracted. All difference heatmaps were generated by taking the two original heatmaps and subtracting the signal in each bin. Metaplots were created by averaging the signal across all features for each bin in the corresponding heatmap.

### Motif analysis

Motif enrichment analysis was performed using CentriMo v4.12.0 from the MEME suite. Enrichment was searched for within the regions 1 kb upstream to 1 kb downstream of the centers of the DOCRs. Motifs tested were the “HOCOMOCOv11_core_HUMAN” set obtained from the MEME motif database v12.21. Footprinting analysis was performed using the TOBIAS suite v0.13.1 with the ATACorrect, FootprintScores, and BINDetect tools. Motifs used with TOBIAS were the “JASPAR2022_CORE_vertebrates_non-redundant_pfms” set obtained from the JASPAR database.

### Pathway analysis

DOCR regions were associated with the nearest upstream or downstream gene, sub-divided by DOCR direction (GAIN, LOST) and genomic context (genic, non-genic), generating four sets of genes: GAIN genic, GAIN nongenic, LOST genic, LOST nongenic. Each gene set was tested for hypergeometric enrichment using clusterProfiler-4.6.2 (37) with gene sets defined by MSigDB obtained using msigdbr-7.5.1 (https://CRAN.R-project.org/package=msigdbr). Enrichments were tested separately using Gene Ontology Biological Process (gobp), Gene Target Regulatory Data (GTRD), and Hallmark Pathways. Pathways with adjusted P-value 0.05 and at least four genes were used for multienrichment analysis with R Github “jmw86069/multienrichjam” (38). The four sets of pathways were integrated to form a pathway-gene incidence matrix, from which four functional clusters were defined using hierarchical clustering. Pathway clusters and genes were represented as a concept network, color-coded by input gene set. Genes with significant expression changes by RNA-seq were indicated with a colored border around each gene node.

### Biological relevance of MG132 DOCRs

Processed ATAC-seq coverage files were downloaded for 74 TGCA BRCA tumor samples (39). Signal from technical replicates was merged for each sample using unionBedGraphs from the BEDTools suite.

### TCGA Breast Cancer ATAC Signal Heatmaps

Signal coverages were summarized for each TCGA sample, at DOCR positions which overlapped the called super-enhancers. Samples were split into Basal or Non-Basal sub-groups (39). Data were visualized using ComplexHeatmap (40) with sample columns, and super-enhancer DOCRs rows, displaying row-centered signal. Samples were split into Basal or Non-Basal sub-groups, and rows were split into six sub-clusters after hierarchical clustering.

### ER ChIP-seq Signal Heatmaps

ER ChIP-seq signals from (41) were summarized at DOCRs overlapping called super-enhancers, by quantifying reads across 20 bins scaled evenly across each super-enhancer region, with 2,000 bases upstream and 2,000 bases downstream the super-enhancer region each divided into ten 200 base bins. ER ChIP-seq signals were adjusted by subtracting input signal from ER-IP signal within each treatment group, estradiol (E2) or vehicle. Adjusted signals were displayed as heatmaps, with the same row ordering and sub-clusters defined by the TCGA breast cancer heatmaps.

### Start Sequencing data processing

Raw reads were quality filtered to include only those with a mean Phred quality score of 20 or greater. Adapter was trimmed using Cutadapt version 1.12 (42). The preprocessed reads were aligned to the hg19 assembly using Bowtie version 1.2 and parameters -v 2 -m 1 --best –strata (33). TSS calls were assigned from the aligned reads using GENCODE v27 and the TSScall tool. TSScall is based on methods used in previous studies (43), and it is available on GitHub (https://github.com/lavenderca/TSScall). In addition to identifying TSS locations, this tool also categorized the TSS calls as either associated with an annotated TSS (obsTSS) or not associated with an annotated TSS (uTSS), and it identified whether the TSS calls have a downstream antisense (divergent) TSS. This information was used to further categorize the obsTSSs as either having a divergent TSS that was also another obsTSS (bidirectional) or having a divergent TSS that was a uTSS (PROMPT). Also, any uTSS that overlapped any part of a gene body was further categorized as genic, whereas all other uTSSs were categorized as intergenic.

## RESULTS

### Changes in chromatin accessibility in response to MG132 treatment

Chromatin accessibility was assessed by Assay for Transposase-Accessible Chromatin with high-throughput sequencing (ATAC-seq) (29) after 0, 4, and 24 hour treatment with MG132 resulting in a total of 162,546 merged peaks from which a subset of 14,008 high confidence OCRs were tested for differential accessibility **(Figure 1, Figure S1A-E, Table S1)**.

**Figure. 1.**
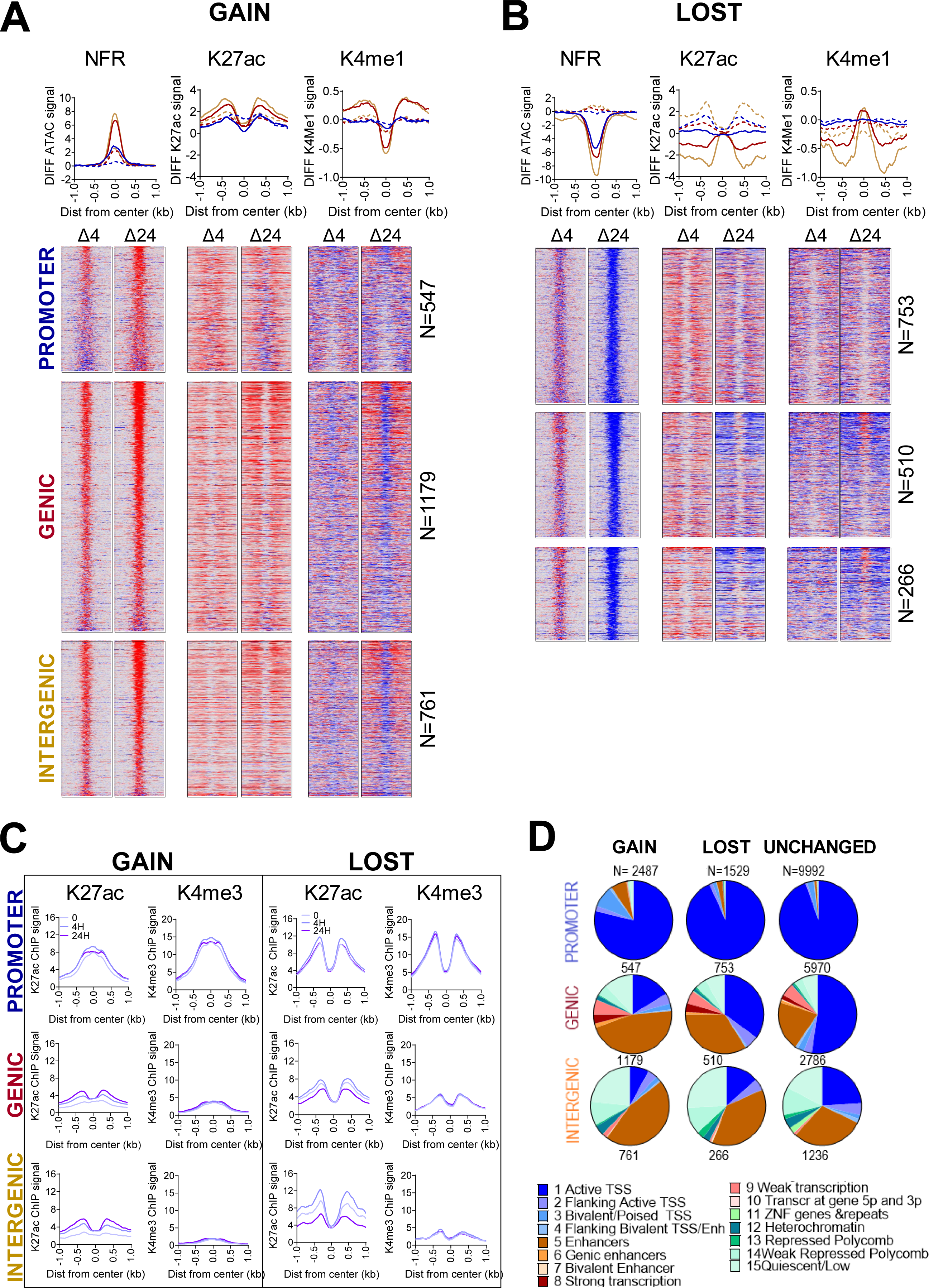
Changes in chromatin accessibility following MG132 treatment. **A)** Heatmaps (bottom) and metaplots (top) showing differential (MG treated minus untreated) signal for DNA accessibility (Nucleosome free reads, NFR), H3K27ac and H3K4me1 at DOCRs that <u>increase (GAIN)</u> and **B)** <u>decrease (LOST)</u> accessibility after 24H treatment compared to untreated. Signal spans ±1kb from the center of the defined DOCR regions and is ranked based on the degree of change in accessibility, where regions at the top have the most change in accessibility. The color scale shows an increase (red) or decrease (blue) in differential signal. Side by side heatmaps show differential signal at the 4 and 24H time points. DOCRs are split by genomic category into **PROMOTER**, **GENIC**, and **INTERGENIC**. The dashed lines in the meta plots correspond to differential signal at 4H, and solid lines at 24H. N is the number of DOCRs in each genomic category. **C)** Metaplots for ChIP-seq binding signals of H3K27ac and H3K4me3 at DOCRs. Signal includes the untreated state (0) and after 4H and 24H of treatment. **D)** Pie charts showing the proportion of DOCRs assigned to various chromatin states using Epilogos https://epilogos.altius.org/. N is the number of DOCRs in each category.

Of the 14,008 OCRs tested, 346 and 4,016 were differentially, increased or decreased, open regions (DOCRs) in cells treated with MG132 for 4 and 24 hours, respectively **(Table S1).** The size distribution of open chromatin regions was similar for unchanged and differential regions, suggesting that the size of the regions was not the key factor determining the differences in DNA accessibility **(Figure S1D)**. The distribution of distances to nearby TSSs of the ATAC peaks did not differ between treatment conditions **(Figure S1E)**. However, while 65% of the aggregate tested OCRs were close to a TSS (<±5kb), only 10% (4H) and 40% 24H) of the DOCRs were close to a TSS (**Figure S1F**). Further analysis of the genomic features revealed that regions with increased accessibility were predominantly in genic and intergenic space, while a higher percentage of those with decreased accessibility were promoter proximal (-1 to +0.5 kb) (**Figure S1G**). Additionally, although there was a slight bias toward DOCRs with increased accessibility (GAIN) across chromosomes, three chromosomes (Chr 16, 19 & 22) were primarily enriched for DOCRs with decreased accessibility (LOST) (**Figure S1H**).

### Chromatin landscape of MG132 DOCRs

There was a time-dependent change in accessibility following MG132 treatment, with most of the changes occurring in cells treated with MG132 for 24hours. Also, regions that showed an increase in accessibility during the 4-hour treatment largely overlapped with those that increased accessibility during the 24-hour treatment **(Figure S1I)**. For this reason, downstream analysis was performed with the 4,016 regions, of which 2,487 were more accessible (GAIN) and 1,529 were less accessible (LOST) in cells treated for 24H compared to control (**Table S1, Figure S1J**). Regions that became more accessible showed a progressive increase in DNA accessibility from 4H to 24H (**Figure S1J**). In contrast, the less accessible regions, show little decrease at 4H and don’t appear to significantly decrease until 24H. (**Figure S1J**). This result suggests that genomic regions that become accessible following MG132 start to do so rapidly (within 4H), whereas regions that become less accessible start to compact later.

To understand whether the differences in accessibility depend on their genomic context, we classified the DOCRs based on their genomic space: proximal to a TSS (promoter), within the transcriptional unit of a gene (genic), or outside of a gene (intergenic) (**Figure S1G**). The GAIN DOCRs showed a larger magnitude of change in the genic and intergenic space compared to the promoter proximal regions [**(Figure 1A)**, compare NFR enrichment shown on metaplot]. In contrast, in the LOST DOCRs, all three genomic regions show a similar decrease in DNA accessibility **(Figure 1B)**. Additionally, at 24H, promoter proximal regions that increase accessibility do not reach the same level of openness as of those that decrease accessibility in untreated cells [**(Figure S1N),** compare NFR GAIN 24H to NFR LOST 0H]. Thus, MG132 treatment results in both decompaction and compaction of chromatin, where the degree of change in accessibility is influenced by the genomic context, leading to reprogramming of accessible sites in the genome.

### DOCRs represent distinct cis-regulatory elements

DNA accessibility is regulated through the actions of multiple histone post-translational modifications (5). Some well characterized post-translational modifications of histones colocalize with specific chromatin states. Since a majority of DOCRs were distal to TSS, this suggests that many DOCRs could be cis-regulatory elements. To determine whether DOCRs were enriched for specific cis-regulatory elements, we performed ChIP-seq with antibodies against, H3K4me3, together withH3K27ac and H3K4me1, histone marks, generally enriched at promoter and enhancer elements, respectively (44,45) **(Figure 1A, B, & C)**.

The H3K27ac mark was differentially associated with both GAIN and LOST DOCRs. Promoters were generally enriched with H3K27ac and H3K4me3, but largely depleted of H3K4me1 (**Figure 1**). Further, promoter regions with increased accessibility showed a distinct chromatin architecture compared to those with decreased accessibility, based on differences in the shape of the H3K27ac and H3K4me3 signal (**Figure 1C**). In regions that increase accessibility, H3K27ac and H3K4me3 accumulate at the center of the accessible region, exhibiting a unimodal profile. However, in regions that decrease accessibility, both marks flank the center of the region, in a bimodal profile, which is generally associated with canonical promoters **(Figure 1C & Figure S1L & M)**. Analysis of the enrichment plots derived from mono-nucleosome fragments, total H3, H3.3 and nucleosome free fragments, also reveal distinct differences between the two classes of promoters (**Figure S1N**). At the promoters with increased accessibility, the mono nucleosome signal is enriched at the center of the region but depleted in regions with decreased accessibility. Compared to LOST, GAIN promoters show higher total H3 and lower H3.3 signal though enriched with H3K122ac at center of the region. Taken together, these results suggest distinctive chromatin architectures at promoters that increase compared to those that decrease DNA accessibility following MG132 treatment.

DOCRs at genic and intergenic regions were also differentially enriched with histone marks. Overall, the largest changes in H3K27ac were observed in genic and intergenic regions, where H3K27ac was enriched and depleted at regions that increase and decrease accessibility, respectively **(Figure 1)**. Similarly, genic, and intergenic regions were also enriched or depleted of H3K4me1, indicating gain and loss of enhancer activity following treatment. Further, for H3K27ac, genic and intergenic regions always predominantly display bimodal profiles of the H3K27ac, irrespective of DOCR class **(Figure 1C)**. Genome browser shots represent examples of genomic loci showing distinct differences in the chromatin landscape of DOCR classes **(Figure S1O)**.

We also explored the relationship of DOCR directionality with genomic region and Epilogos chromatin state data (http://epilogos.altius.org/) (36) **Figure 1D**. Promoter regions were largely associated with the transcriptional regulatory function of active TSS **(state 1),** but there were also differences in chromatin architecture between GAIN and LOST promoters. GAIN promoters showed a higher enrichment of states marked by bivalent/poised TSS and enhancer states (**state 3 &5),** whereas LOST promoters look more like unchanged promoters (**Figure 1D)**.

Further analysis of chromatin states also revealed additional inherent differences in chromatin architecture of genomic regions affected by MG132. Genic regions with increased accessibility were associated with fewer transcriptionally active states (**state 1-4**) compared to regions with decreased accessibility, but both DOCR classes overlapped proportionally more enhancer regions than unchanged regions (**state 5-7**) **Figure 1D**. DOCRs in intergenic space were associated with enhancer states (**states 5-7**) more so than unchanged regions, while all three classes of intergenic OCRs showed association with repressive chromatin states (**states 11-14**) **Figure 1D**. In summary, integrating ATAC-seq and ChIP-seq of histone marks revealed distinct chromatin architecture of promoter DOCRs and uncovered a subset of genic and intergenic regions associated with putative active and poised enhancers in MG132 treated cells.

### DOCRs are sites of RNAPII transcription

Chromatin impedes transcription and the dynamic changes in chromatin accessibility that we observe in MG132 treated cells, may have potential consequences on transcription. We next performed ChIP-seq with antibodies against RNAPII, to profile the genome-wide occupancy of RNAPII as a proxy for monitoring transcribed regions in MG132 treated cells. Regions that increase accessibility upon treatment were enriched with RNAPII, where the largest changes in Pol II binding occurred at promoter and to a lesser degree at intergenic DOCRs (**Figure 2A**). Regions with decreased accessibility exhibited different Pol II binding patterns depending on the genomic region. Promoter regions with decreased accessibility were highly enriched with non-phosphorylated Pol II (Non-Phos) compared to serine 5 phosphorylated Pol II (Ser5-P) (**Figure 2B**). In contrast, genic and intergenic regions with decreased accessibility were largely depleted of Ser5P, while some regions retained non-Phos RNAPII binding. There were also distinct differences in RNAPII profiles between the regions that increased compared to those that decreased accessibility (**Figure 2C**). At promoters and genic regions with decreased accessibility, RNAPII signal shows a bimodal profile, in contrast to the sharp peak observed at regions with increased accessibility.

**Figure 2.**
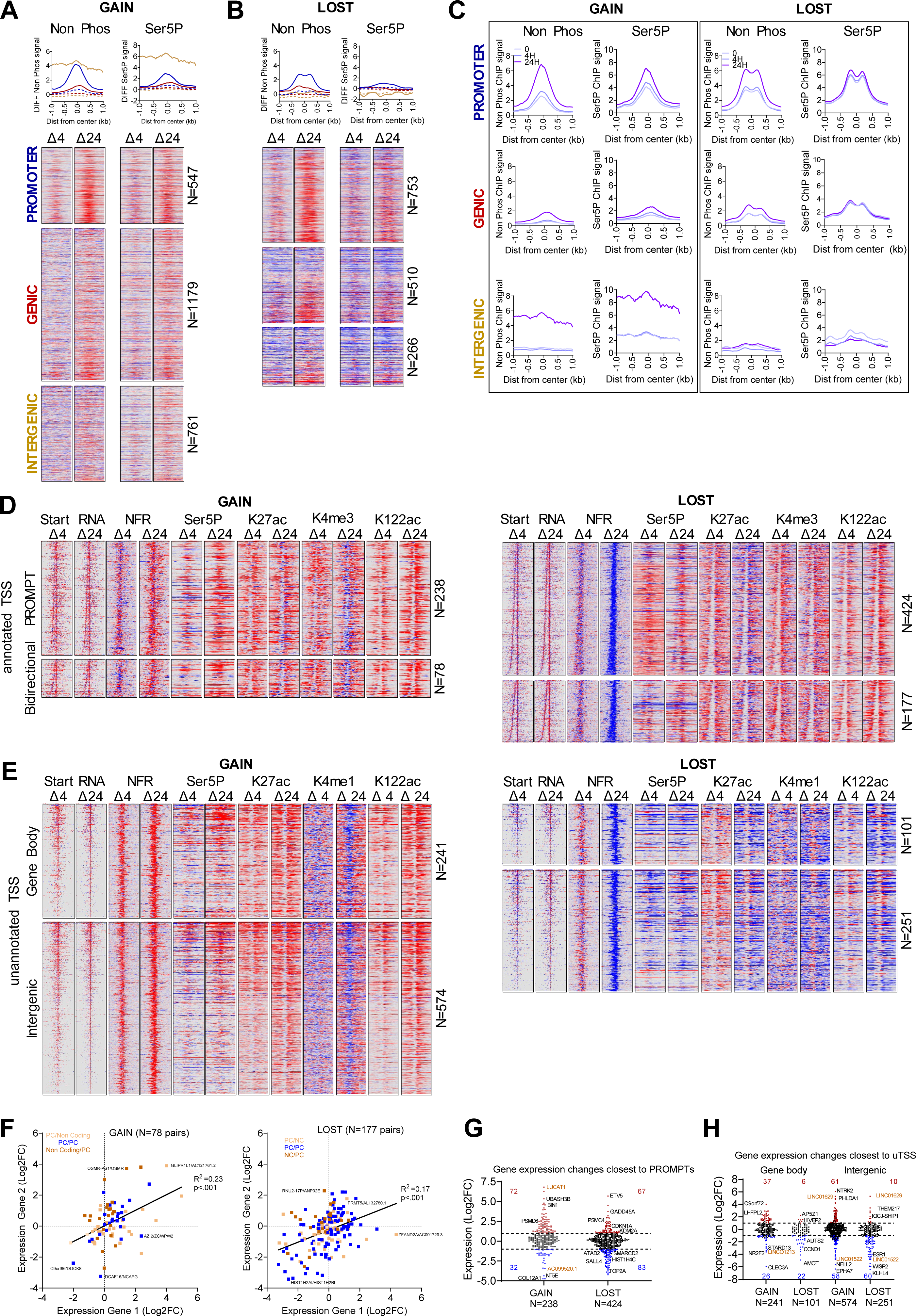
Differential open chromatin regions are sites of RNAPII transcription. **A)** Heatmaps (bottom) and metaplots (top) showing differential signal for Non-Phosphorylated and Ser5P RNAPII at DOCRs that increase and **B)** <u>decrease</u> accessibility. Signal spans ±1kb from the center of the DOCRs and is ranked based on the degree of change in accessibility. **C)** Metaplots for ChIP-seq binding signals of Non-Phosphorylated and Ser5P RNAPII at DOCRs. Signal includes the untreated state (0) and after 4H and 24H of treatment. **D)** Heatmaps showing differential signal for start RNA, accessibility (NFR), RNAPII (Ser5P), and active histone marks at genic TSSs with divergent transcription that overlap GAIN (left) and LOST DOCRs (right). Signal spans ±500 bases from the TSSs and is ranked by distance between sense and antisense TSS pairs, where TSSs at the top have the largest distance. TSSs are split by category into Promoter Upstream Transcripts (PROMPT) and Bidirectional (head-to-head) TSS pairs. N is the number of TSS pairs in each category. **E)** Heatmaps showing differential signal as in Figure 2D, but at nongenic TSSs that overlap DOCRs. TSSs are split by genomic category into gene body and intergenic. **F)** Scatter plot showing gene expression changes of bidirectional (head-to-head) TSS pairs. TSS pairs are colored by gene class, where each TSS in the pair is either protein coding (PC) or non-coding (NC). **G)** Violin plot showing gene expression changes of genic TSSs associated with promoter upstream transcript (PROMPT) category. Significantly different genes (FDR≤0.05, Log2FC±1) are colored red (upregulated) and blue (downregulated). Genes labeled in gold are noncoding genes. **H)** Violin plot showing gene expression changes of genes closest to nongenic TSSs, which are split by genomic category into gene body and intergenic

### DOCRs are sites of pervasive transcription

Since RNAPII is enriched at DOCRs, we next determined the level of transcription from these regions. We found most regions had detectable RNA synthesis based on RNA-seq read counts in the region spanning +/- 250 bp from the center of each DOCR **(Figure S2A).** RNA counts showed small increases in expression at regions with increased accessibility, while intergenic regions with decreased accessibility had substantial decrease.

To evaluate further how the changes in accessibility affected transcription initiation, we performed start-seq to assess enrichment of transcription initiation start sites at DOCR classes. We defined the TSSs as either being associated with a known gene (genic), or not associated with a known gene (non-genic), then identified genic and non-genic TSSs that overlapped with GAIN or LOST DOCRs.

We found 316 and 601 genic TSSs that overlapped with GAIN or LOST DOCRs, respectively. Since these primarily represent start sites from promoter regions, which we found to differ in chromatin architecture, we tested whether there were differences in patterns of transcription initiation **(Figure S2B, Table S2)**. For this, we first identified genic TSSs that were paired with an upstream divergent TSS. Then, we categorized the genic TSSs based on whether its divergent pair was non-genic TSS, making it an annotated mRNA gene associated with an unstable antisense transcript [promoter upstream transcript (PROMPT)], or whether its divergent pair was another genic TSS, making it a bidirectionally active TSS of mRNA-mRNA pairs arranged head-to-head. We found 70% of the divergent transcription from promoters that increase (GAIN) and decrease (LOST) accessibility was due to PROMPTs, whereas 30% was due to bidirectional (head-head) mRNA gene pairs (**Figure 2D**).

To explore this further, we sorted the PROMPTs and head-to-head pairs based on genomic distance between the sense and antisense TSS pairs, and evaluated the relevant start-seq, RNAPII binding, and selective histone modifications. This analysis revealed a striking contrast in the distance between divergent TSSs from DOCRs GAIN compared to LOST (**Figure 2D, start-RNA**). The median distance between the aggregate TSS pairs was wider in regions of decreased accessibility (277 nt) vs increased accessibility (188 nt) accessibility [(p<0.0001) **(Figure S2C)]** and this difference in divergent distance persisted when we stratified TSS into PROMPTs and head-to-head pairs **(Figure S2D).** Heatmap profiles representing ChIP signals for active histone marks and RNAPII (Ser5P) around the TSS sites reflected the differences in chromatin architecture and the divergent transcription patterns occurring at the two promoter classes (**Figure 2D**).

We also explored non-genic TSSs that overlapped with DOCRs **(Table S3)**. We found more non-genic TSSs in regions that increase (815) compared to those that decrease (352) accessibility (**Figure S2E**), which we subsequently divided into TSS sites initiating from either gene body or intergenic regions. Over 70% of non-genic TSSs were intergenic and 30% were in the gene body in regions of increased (GAIN) and decreased (LOST) accessibility, respectively (**Figure 2E**). As shown on the heatmaps, regions of increased accessibility were enriched with start RNA, RNAPII, and active enhancer marks, while these features were depleted in regions with decreased accessibility. Thus, proteasome inhibition had a large impact on transcription initiation patterns, which were largely influenced by chromatin architecture resulting from MG132 treatment.

We next used RNA-seq to test the effect of transcription initiation patterns on gene expression. Bidirectional gene pair expression showed a slight positive correlation in regions that both increase and decrease accessibility (r^2^ =0.23 and r^2^ =0.17), and some of the genes (46 and 108) were differentially expressed [36 UP vs 10 DN and 40 UP vs 68 DN (FDR <0.05; Log2FC =1) **Figure 2F and Table S2)]**. In regions with increased accessibility, more than 65% of the bidirectional genes were composed of protein coding (PC) and non-coding (NC) pairs, whereas only 42% of the pairs were PC/NC in the regions with decreased accessibility. Upregulated genes were typically chromatin regulators and down regulated genes were predominantly histone H2A and H2B transcripts, examples of which are highlighted **(Table S2).**

We observed that in both GAIN and LOST regions 85-95% of annotated TSSs associated with a PROMPT were protein coding genes (**Figure 2G**, **Table S2)**. However, gene expression changes related to accessibility were different. Genes in GAIN regions were generally induced (72) compared to those repressed (32), whereas genes in the LOST regions were similarly induced (67) and repressed (83) by MG132 [(FDR <0.05, Log2FC =1) (**Figure 2G**, **Table S2)]**. These results are consistent with distinct effects of upstream antisense transcription (PROMPT) on host gene expression. Examples of induced and repressed genes for both classes of accessible regions are highlighted (**Figure 2G)** and browser tracks present chromatin state and transcription initiation sites associated with examples of genes expressed from each class of TSS pairs (**Figure S2F**).

### Genic and Intergenic DOCR transcription is largely noncoding

Having characterized transcription events occurring at promoter regions affected by MG132, we next explored transcription occurring at regions distal to TSSs. In non-genic regions with increased accessibility, Ser5-P, H3K4me1, H3K27ac, and H3K122ac were enriched, and they were depleted in regions with decreased accessibility, inferring actively transcribed cis-regulatory elements (**Figure 1A &B and Figure 2E**). The degree of acetylation and chromatin accessibility in the coding regions was associated with aberrant activation or repression of unannotated promoters as shown by start RNA expression (**Figure 2E**).

We explored whether non-genic RNAPII initiation had any effect on gene expression changes in nearby genes (**Figures 2H,S2G**). Regions with increased accessibility had a relatively similar number of up and down-regulated genes nearby, whereas regions with decreased accessibility had a substantially higher percentage of down-regulated genes. Some examples of induced and repressed genes for both classes of accessible regions are highlighted (**Figure 2H, Table S3)** and genome browser shots present chromatin state and transcription initiation sites associated with specific gene examples **(Figure S2I)**.

Finally, analysis of the start-seq data revealed additional transcription initiation from genic and intergenic regions of cells treated with MG132, results in expression of non-coding RNAs, most of which were uncharacterized novel transcripts whose expression was not detected by RNA-seq **(Figure S2H)**, and those detected were generally repressed **(Figure 2H, Table S3).**

### MG132 DOCRs are enriched with sequence motifs of TF that regulate chromatin and oncogenic factors

In general, sequence specific transcription factors bind cis-regulatory DNA elements within accessible chromatin to orchestrate RNAPII transcription. We applied motif discovery to search for candidate transcription factors (TF) motifs that were enriched at DOCRs that increased and decreased accessibility.

We found distinct classes of TF motifs enriched in regions that gained accessibility compared to regions that are less accessible in MG132 treated cells (**Figure 3A**). AP-1 (FOS/JUN) motifs were notably enriched in genic and intergenic regions that gained accessibility compared to regions that were less accessible. On the other hand, NFY motifs were specifically enriched at less accessible promoters. Motifs for SP and Krüppel-like transcription factors were enriched at all regions, with the greatest enrichment at promoters with decreased accessibility. Finally, Fork head transcription factor motifs were enriched in the genic and intergenic regions, whereas CTCFL (BORIS/CTCF like) motifs were enriched overall but depleted at intergenic regions with decreased accessibility. Transcripts of most TFs associated with significant motifs were expressed in MCF-7 cells (**Figure 3B**). Foot printing analysis of DNA sequences underlying the 14,008 global open chromatin regions using TOBIAS (46) found 267 enriched and 54 depleted motifs, upon 24-hour treatment with MG132 [(pvalue -log10 ≥100 cutoff), **Figure 3C)]**. These analyses confirmed enrichment of the FOS-JUN family of motifs, and depletion of KLF and NFY motifs. TOBIAS also detected additional TF motifs of interest, including enrichment of NFE2 (NFE2L1/2), ATF, MAF, BACH, CEBP, depletion of YY, ARNT, ETS Variant TFs (ETV5 etc.) and potential repressors, including ARID3B/5A, JDP2, SAT1B.

**Figure 3:**
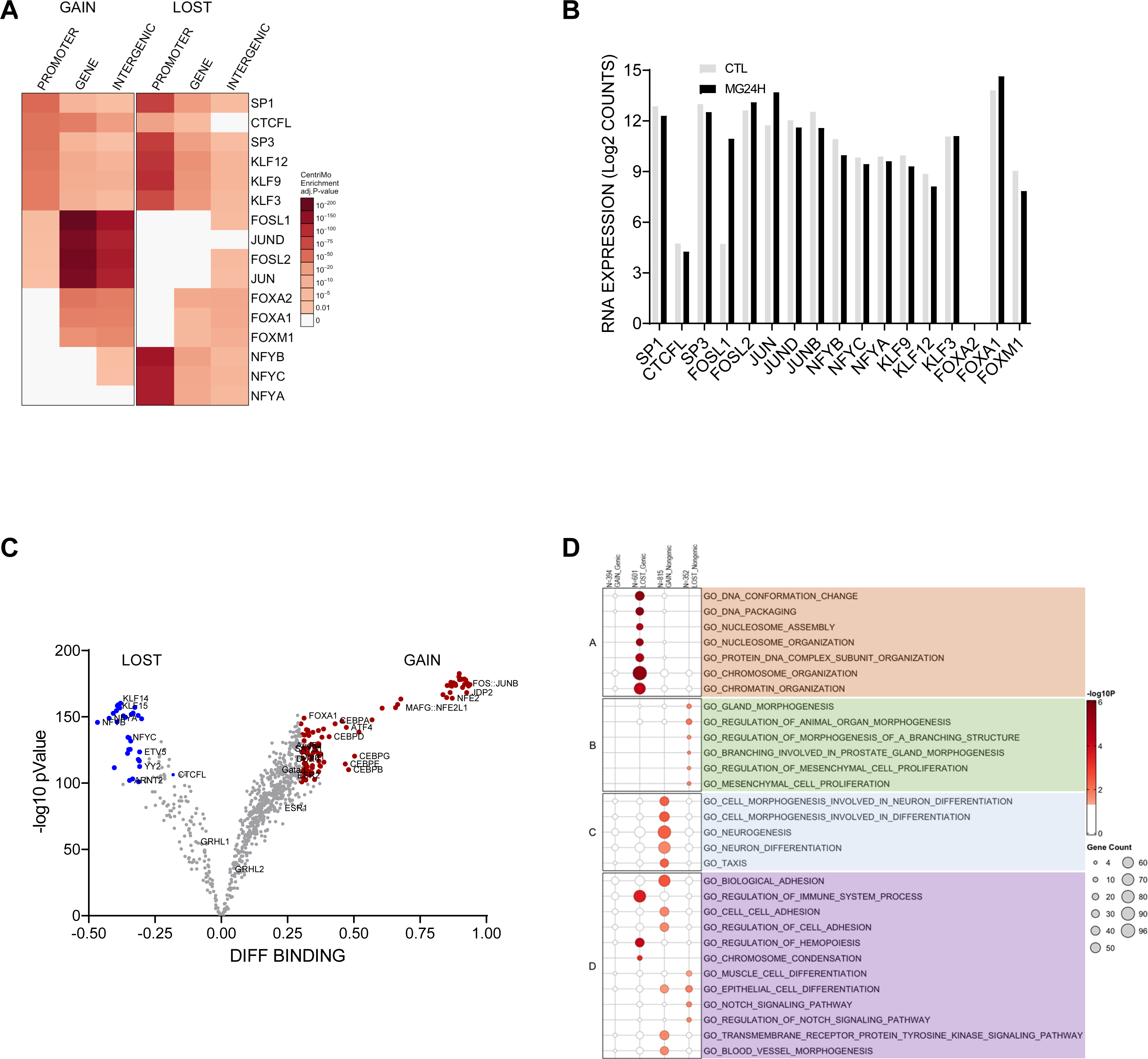
DOCRs are enriched with sequence motifs of TF that regulate chromatin and oncogenic factors. **A)** Heatmap showing the degree of enrichment for top 3 transcription factors binding motifs identified as enriched in each DOCR category. Each row corresponds to a TF. Each column corresponds to the DOCR genomic category and the heatmap is split into DOCR class left (GAIN) and right (LOST). **B)** Relative mRNA expression of TFs enriched at DOCRs, showing RNA-seq read counts in control (CTL) and 24H treated cells. **C)** Volcano plot showing the degree of differential binding and the statistical significance of the difference for all TF motifs queried. TF motifs with significant differential binding are colored red (increase) and blue (decrease). **D)** Dot plot showing the degree of enrichment for gene lists enriched with genes closest to DOCRs split by category. Dot size represents the number of genes in each GO biological processes.

Next we tested for enrichment of Gene Ontology biological processes in the genes nearest to the DOCRs. DOCRs with decreased accessibility in promoter regions were enriched in chromatin organization, nucleosome assembly, and DNA packaging (**Figure 3D, cluster A**). Most genes encoding histones, histone chaperones, and chromatin regulators also appear in cluster A of the concept network (cNET) (**Figure 3SA**). Genes in promoter regions with increased accessibility showed no significant enrichment, perhaps because a majority of the TSS code for noncoding RNAs (**Figure 2F**). Genes whose TSS were close to distal DOCR regions with increased accessibility were enriched for processes involved in neurogenesis (cluster C) and cancer-related processes (cluster D) including cell adhesion. On the other hand, genes in regions of decreased accessibility showed enrichment in processes involved in cell fate and cancer, including gland morphogenesis (cluster B), cell proliferation, and Notch signaling (cluster D). Genes shared by cluster A and D represent a link between chromatin organization and cancer hallmark processes, involving FOXA1, ESRI, PHF14 and the ATPase proteasome subunits, PSMC5 and PSMC4 (**Figure S3A)**.

A follow-up analysis revealed DOCR linked genes were specifically enriched in a few Hallmark pathways encompassing a myriad of pathological processes that may impact breast cancer pathophysiology: mTORC1, Hedgehog signaling, and estrogen response, controlling cell proliferation; E2F regulation control of cell cycle and replication; IL2/STAT5 and ROS signaling involved in ferroptosis cell death and immune response, (**Figure 3SB)**. Representative genes of each node are shown in the cNET plot (**Figure S3C**).

Finally, we utilized Gene Transcription Regulation Database (GTRD; http://gtrd.biouml.org/) to explore additional transcriptional regulators of DOCR genes (47). We find enriched targets of the histone lysine methyltransferase ASH1L, the TF CEBPZ, and PRKDC (DNA-PK) associated with genes in regions with decreased accessibility (**Figure S3D & S3E**). On the other hand, regions with increased accessibility represent targets of neurogenesis regulators HES2, PAX3, CBX5, PRDM5, and chemoresistance factor MED16 (**Figure 3D & S3E,**).

Taken together, after 24-hour proteasome inhibition, we found increased accessibility in genes associated with neurogenesis, and decreased accessibility in genes associated with chromatin maintenance and cell cycle regulation.

### MG132 DOCRs distinguish breast cancer tumor subtypes

We evaluated the accessibility of MG132 DOCR regions in breast cancer tumors by leveraging data from (39), where they analyzed chromatin accessibility in primary human cancers including breast cancer. We plotted accessibility across breast cancer samples for DOCRs, subdivided by promoter, genic, and intergenic regions **(Figure 4A)**. The hierarchical clustering of open chromatin signal within DOCRs clearly separated non-basal (hormone receptor/estrogen receptor (ER) positive) from basal (hormone receptor negative) breast cancer subtypes, with differences more substantial in genic and intergenic compared to promoter DOCRs (**Figure 4A**).

**Figure 4.**
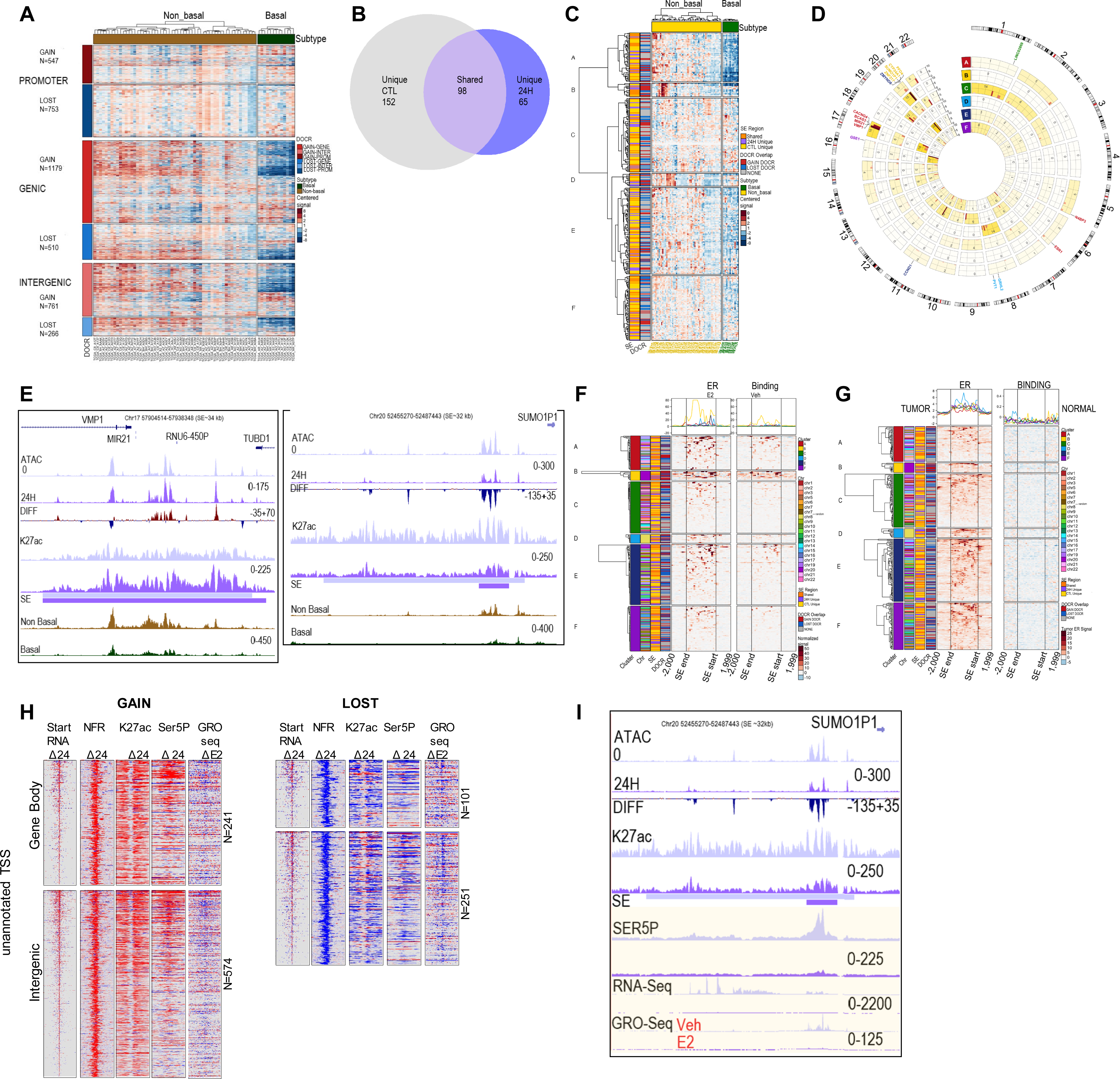
DOCR regulatory elements are enriched in non-basal breast cancer tumors. **A)** Heatmap showing chromatin accessibility signal (ATAC-seq) of TCGA breast cancer tumors (1) in DOCRs. Each row corresponds to a DOCR, and they are split by increase (GAIN) or decrease (LOST) in accessibility and by genomic category (PROMOTER, GENIC, INTERGENIC). N is the number of DOCRs in each category. Each column corresponds to a TCGA breast cancer tumor, split by non-basal (N=57) and basal (N=13) subtypes. Signal is z-score normalized by row, where red is high relative signal and blue is low relative signal. **B)** Venn diagram highlighting the number of super enhancers identified in Control only, 24H treated only, or shared between the samples. **C)** Heatmap showing chromatin accessibility signal (ATAC-seq) of TCGA breast cancer tumors in SEs. Each row (N=333) corresponds to a SE region, and they are split by cluster from hierarchical clustering. **D)** Circos plot showing chromosome coordinates of SEs, split by cluster. The outer ring shows reference chromosomes 1 through 22 in clockwise orientation. Inside rings correspond to clusters A-F from the heatmap in Figure 4C. The number of SE regions in each cluster per chromosome is indicated inset. Representative genes in SE regions in each cluster are shown. Color density indicates SE enrichment within each cluster and chromosome. **E)** Browser tracks showing the VMP1/MIR21 and SUMO1P1 super enhancer regions. Tracks show read coverage of chromatin accessibility (ATAC-NFR), differential read coverage of accessibility (DIFF), H3K27ac read coverage, super enhancer regions (SE), and average ATAC read coverage for non-basal (brown) and basal (green) TCGA breast cancer tumors. Each track represents a control (0) light and 24H sample dark purple **F)** Heatmap showing ER ChIP-seq signal of MCF-7 cells treated with E2 (left) or Vehicle (right) in SE regions. Signal spans from 2kb upstream of the super enhancer, through the body of the super enhancer, and to 2kb downstream of the super enhancer. Rows are split by cluster from the heatmap in Figure 4C. Signal is z-score normalized by row, where red is high relative signal and blue is low relative signal. **G)** Heatmap showing ER ChIP-seq signal of breast tumor (left) or normal tissue (right) in SE regions. **H)** Heatmaps showing differential signal for start RNA, accessibility (NFR), RNAPII (Ser5P), K27ac and GRO-seq (E2) at nongenic TSSs that overlap GAIN (left) and LOST DOCRs (right). **I)** Browser tracks showing the SUMO1P1 super enhancer region. Tracks show read coverage of chromatin accessibility (NFR), differential accessibility (DIFF), SE regions, K27ac, RNAPII (SER5P), RNA-seq, and GRO-seq from cells treated with veh and E2.

### SE elements enriched with DOCRs are predominantly accessible in non-basal breast cancer tumors

We next defined super enhancers (SE) using ROSE, which identified 250 in control and 164 in MG132 treated cells (**Figure S4A, Table S4)**. Among these, 98 SEs were shared between control and MG132 treated cells, and most of them (82 SEs, 84%) overlapped with at least one DOCR (**Figure 4B, Table S4**). Most (∼65%) SEs unique to control cells did not overlap DOCRs, whereas SEs unique to MG132 treated cells showed higher overlap with DOCRs, and these were primarily in regions that increased chromatin accessibility (**Figure 4B, Table S4**). Additionally, SEs unique to MG132 treatment had lower H3K27ac signal compared to the shared SEs (**Figure S4B**). Representative examples of each class of SE are shown (**Figure S4C**).

Based on this analysis, we sought to understand the biological significance of the SEs in breast cancer biology. We examined the TCGA breast cancer ATAC-seq signal at the SE regions which overlap open chromatin regions **(Figure 4C, Table S5)**. Hierarchical clustering was used to split the SEs into 6 sub-clusters with distinct DNA accessibility profiles. Cluster A contained 54 SEs which showed substantially lower accessibility in basal compared to non-basal breast tumors. Interestingly, 16 cluster A SEs were located on chromosome 17 (**Figure 4D**). One such SE which overlaps the VMP1/MIR21 gene locus showed increased accessibility with MG132 but decreased accessibility in basal tumors (**Figure 4E & S4D**). Other SEs of interest in cluster A were in regions that show decreased accessibility in treated cells, including an SE in the BCAS3 intron, an SE in nearby of CACNG4, a region in the proximity (∼13kb) of N4BP3 (Chr 5) and ESR1 (Chr 6), (**Figure 4D, highlighted, & Table S5**).

Clusters B and D consisted of 15 and 14 SE regions, respectively, and showed high accessibility in a subset of non-basal breast tumors. All cluster B SE regions were located on chromosome 20, and all cluster D regions were on chromosome 8 (**Figures 4C & 4D)**. While accessibility in basal regions of both clusters B and D was low, the degree of accessibility in cluster B regions on chromosome 20 was even lower than cluster D. The majority of cluster B SEs (14/15) overlapped with DOCRs, and 10 were in regions of increased DNA accessibility. The regions that decreased chromatin accessibility are in Chr 20 gene deserts, neighboring non-coding RNAs. One such region is near the SUMO1P1 pseudogene and shows a dramatic decrease in accessibility in MG132 treated cells, along with high accessibility in non-basal breast tumors (**Figure 4E, right panel).** Further, the SUMO1P1 desert region is more accessible in highly proliferative compared to lowly proliferative breast tumors (**Figure S4D**). Cluster D SEs also largely overlapped DOCRs (11/14) and included SEs nearby PVT1 and GRHL2 **(Figure 4D, Table S5).**

Cluster C, E and F SEs showed modest changes in accessibility, but differences across breast tumors were still evident (**Figure 4C**).

Summary figures for SE regions near VMP1/MIR21 (Chr 17), SUMO1P1 (Chr 20), and PVT1 (Chr 8) are shown as representative examples from cluster A, B, and D, respectively, showing higher accessibility in non-basal compared to basal breast sample tumors **(Figure S4D).** Figures depicting SE regions near LINC02869 (Chr1), ZMYND8 (Chr 20), and GSE1 (Chr 16) are shown as representative examples of clusters C, E, and F, respectively **(Figure S4E)**.

Non-basal breast tumors are characterized by the expression of estrogen receptor (ER), so we investigated whether the SE regions were associated with ER binding sites **(Figure 4F & 4G).** To this end, we downloaded publicly available ChIP-seq data for estrogen receptor (ER) in MCF-7 cells and breast tumor samples (41) to examine ER binding at SE regions. We found enriched ER binding in MCF-7 cells concentrated in clusters A and B, which were clusters highly accessible in non-basal compared to basal tumors (**Figure 4F**). We also found these regions were enriched with ER in ER-positive tumors compared to normal breast tissue **(Figure 4G).**

Cancer cells are highly dependent on an active proteasome, to modulate gene regulatory networks that drive cell growth, an important question is whether MG132 represses aberrant transcription associated with ER dependent transcription. DOCRs in the gene body and intergenic regions are transcriptionally active in MG132 treated cells **(Figure 2E).** Next we leveraged data from GRO-seq (48) of ER regulated transcription in MCF-7 cells, to examine whether transcription at these regions was sensitive to E2 treatment. We observed transcriptional induction and repression at sites with both increased and decreased accessibility upon E2 treatment as shown on heatmaps of GRO-seq signal detected across non-genic TSS overlapping DOCRs in cells treated with vehicle or estradiol (E2) **(Figure 4H & S4F)** We then focused on a region in Cluster B containing intergenic DOCRs with decreased accessibility on chromosome 20 (chr20:52462606-52487433), which also includes a large SE that overlaps DOCRs close to SUMO1P1 and shows increased ATAC signal in proliferative non-basal tumors (**Figure 4I, S4D, center panel)**. In MG132 treated cells, the region showed a consistent decrease in accessibility, RNAPII binding, and RNA transcription (**Figure 4I**). Based on GRO-seq, compared with vehicle (veh), E2 treatment also repressed transcription from this region. The decrease in RNA expression seen in the RNA and GRO-seq was confirmed by qPCR using primers spanning the SE region (**Figure S4G**) and the results further validated using known ER targets, TFF1, ZNF217 **(Figure S4H)**.

These data indicate that proper proteasome function is important for breast cancer biology, as proteasome inhibition partly disrupts ER-dependent transcriptional programs that regulate cellular processes critical for cell growth.

## DISCUSSION

Inhibiting proteasome activity is widely believed to disrupt RNAPII transcription initiation, ceasing global gene transcription and expression, although accompanying chromatin changes leading to these events are largely speculative and unexplored (7). Using a low dose of small molecule MG132 enables investigation of proteasome inhibition effects on chromatin accessibility and RNAPII transcription in breast cancer cells. Promoter DNA accessibility is very sensitive to proteasome inhibition, 50% of the regions that decrease accessibility are in promoter regions, agreeing with existing ideas indicating proteasome activity is required for turnover and recycling of RNAPII preinitiation complexes to allow multiple rounds of transcription [reviewed in (10,49)]. Thus, stalled or poised ubiquitinated RNAPII, evidenced by enriched non-phosphorylated Pol II at these promoters could result in the observed decrease in accessibility (32,50). Additionally, the decrease in accessibility is a result of decreased histone acetylation, or eviction of other histone marks generally enriched at promoters, such H2A.Z (51). Interestingly, a subset of promoters does not require proteasome activity to open and this correlates with acetylated H3K27ac, and H3K122ac together with transcriptionally active RNAPII-Ser5-P whose binding is coincident with accessible chromatin (52-54).

A major finding from this study is the observed distinct differences in the shape of the GAIN and LOST promoters, which present as unimodal and bimodal profiles of histone marks and RNAPII. Previous studies have shown bimodal shaped gene promoters present large nucleosome depleted regions which are associated with widespread divergent transcription (43,55,56). Our study presents additional evidence to show bimodal and unimodal promoters exhibit distinct types of divergent transcription patterns in cells treated with proteasome inhibitor. Taking advantage of high-resolution strand-specific mapping of transcription start sites, we stratify divergent transcription into head-head (mRNA-mRNA) and promoter upstream antisense transcription (PROMPT), to show divergent transcription initiation patterns differ between the two types of promoters, resulting in transcription of distinct classes of genes. Strikingly, at bimodal promoters, bidirectional head-head transcription initiation was predominantly from histone gene cluster TSS pairs, whereas PROMPT-mRNA transcription significantly encoded chromatin regulators and histone chaperones, an observation consistent with studies examining the impact of divergent transcription on the human genome (57-59). In contrast, to bimodal, unimodal promoters, head-head, and PROMPT divergent TSS pairs included a combination of long non-coding RNAs, and protein coding genes not typically expressed in breast cancer cells. These genes were enriched in neural pathways including neurogenesis (60,61). This finding is consistent with a requirement for proteasome function to maintain transcription fidelity and cell/tissue specific gene expression [reviewed in (10)]; (62).

The effects of proteasome inhibition on chromatin accessibility were not limited to promoters. We observed large changes in DNA accessibility at regions distal to TSS which were enriched or depleted of enhancer histone marks and RNAPII-Ser5P, suggesting inhibiting proteasome activity reprograms the enhancer landscape of breast cancer cells. Based on the RNAPII density, the reprogrammed enhancers were actively transcribed as we detected increased and decreased start RNAs in regions where changes in accessibility are observed. Importantly, in contrast to promoters which were less accessible, distal regions were hyperacetylated, and more open, resulting in spurious transcription of non-coding RNAs as detected by start-seq. These observations reinforce the notion that the proteasome is required for transcriptome integrity, and inhibiting its function, results in pervasive transcription, which can result in dysregulation of gene networks that may influence cell fate decisions (62-64).

Indeed, changes in accessibility of the enhancer landscape occur at super enhancer regions that regulate transcription and expression of gene networks associated with cell proliferation and chemoresistance, cell fate decisions critical for breast cancer tumorigenesis (65-68). A subset of the super enhancer regions is predominantly accessible in non-basal compared to basal breast cancer subtypes. Of note these regions are primarily in chromosomes 8,17 and 20 hot spot loci, characterized by genomic amplification, aneuploidy, oncogene translocations, and genomic alterations that result in oncogene activation in breast cancer (69-71). Our findings reveal a yet uncharacterized role for proteasome in regulating the accessibility and activity of cis-regulatory elements, important in breast cancer biology (39,72-74).

Breast cancer subtype classification is generally based on hormone receptor status, where the non-basal and basal subtypes, are estrogen receptor (ER) positive or negative, respectively (75). Estrogen receptors are critical regulators of breast epithelial cell proliferation, a major factor contributing to breast cancer tumorigenesis (76). Gene expression networks controlling cell proliferation were repressed by MG132 treatment supporting proteolytic function of the proteasome in ER-mediated transcriptional activity (24). Concomitantly, here we showed inhibiting proteasome function impacted accessibility of ER bound SE regions in hormone receptor positive breast cancers. Estrogen receptors are ligand-activated transcription factors, whose turnover and transcriptional activity is tightly controlled by the proteasome [reviewed in (77),(78)]. Our findings suggest, MG132 may affect accessibility of ER enriched super enhancers, important in the regulation of ER signaling pathways implicated in breast cancer biology.

Lastly, our study points to chromatin accessibility as a potential mechanism by which proteasome inhibitor drugs exert their anticancer effects. We showed dramatic changes in accessibility and transcription, at a subset of super enhancer regions that distinguish non-basal and basal breast cancer subtypes and are also implicated in regulating expression of genes related to cell proliferation and chemoresistance (39,72-74,79). In addition, MG132 suppresses ER enriched super enhancer transcription, suggesting proteasome inhibition alters the function of a critical transcription factor involved in the biology of hormone receptor positive breast cancers. Super enhancers play vital roles in tumorigenesis and small molecule drugs targeting super enhancer function and the molecular machinery that maintain their chromatin state and transcriptional activation, represent promising therapeutic strategy for cancer treatment (19,80,81). Thus, modulating super enhancer accessibility and transcription, is a potential mechanism by which proteasome inhibitor pharmaceuticals could exert anticancer effects in solid tumors, such as breast cancer. Such a mechanism has been suggested by a recent study showing proteasome inhibition caused H3K27 deacetylation of a super-enhancer in control of transcription of the c-MYC proto-oncogene, which resulted in decreased growth of multiple myeloma cells, a blood cancer where proteasome inhibitors are primarily used as therapy (82). Furthermore, other small molecule chemotherapeutics have been shown to disrupt chromatin accessibility and RNAPII transcription of critical cis-regulatory elements, as potential mechanisms for controlling gene expression networks related to cancer cell death (83,84).

We have shown treatment of MCF-7 breast cancer cells with the proteasome inhibitor MG132 results in differential changes in chromatin accessibility and RNA Pol II transcription. These accessibility changes were discrete such that losses were largely at promoters, and gains were predominantly at regions distal to TSS. Promoters which were affected by MG132, displayed diverse chromatin architecture, resulting in divergent transcription of TSS pairs that code for distinct classes of genes, including the HIST1 histone gene cluster, chromatin regulators and novel non-coding RNAs. MG132 chromatin effects also occurred at regions distal to TSS that overlap super enhancer elements accessible in non-basal compared to basal breast cancer tumor subtypes. Our study reveals, a yet uncharacterized chromatin landscape that may contribute to molecular mechanisms by which disruption of proteasome function affects transcription and expression of gene regulatory networks important in breast cancer biology.

### Limitations of the study

Our study has some limitations. We only use MG132 as the proteasome inhibitor and MCF-7 cells as the representative breast cancer cell line. However, the use of MG132 as an experimental drug to inhibit proteasome activity is largely accepted in the field (85). Furthermore, the biological effects we observe with MG132 are replicated by other proteasome inhibitor drugs (23). MCF-7 breast cancers are routinely used as a model to study hormone receptor positive breast cancers (86), and our findings are relevant to breast cancer biology

The genomics approaches taken in this study, do not identify specific factors responsible for the differences in chromatin accessibility and RNAPII transcription during proteasome inhibition. Nevertheless, we speculate the effects of MG132 on the expression of machinery that influences chromatin architecture, for example histone genes, their chaperones, components of epigenetic enzymes (SMARCD2, KDM2A, ATAD2) and non-coding RNAs can have global implications on chromatin organization, including enhancer reprogramming (65,87-89).

In addition to well characterized chromatin and transcriptional regulators, inhibiting proteasome activity, potentially switches the proteasome from proteolytic to non-proteolytic functions on chromatin. Indeed, we show inhibiting proteasome activity results in a stress-induced negative feedback loop, leading to increased expression of proteasome subunits and particularly, the 19S ATPase subunits (24,90). The 19S ATPase subunits, members of the ATPases associated with various cellular activities family (AAA) (91), are versatile components of chromatin regulators, histone chaperones and RNAPII complexes, and through this feedback loop mechanism, probably hijack some chromatin remodeling activities when proteolytic activity is inhibited (11-13,16). In fact, although majority of the evidence leading to this suggestion was obtained from yeast (64), in mammalian cells, ATPase subunits bind to hormone receptor gene promoters to modulate chromatin and transcription (32,92,93), they are enriched at cell type specific enhancesomes (62,94), and are components of transcriptionally active condensates and RNAPII complexes (11,12,16,95). Incidentally, this moonlighting activity of the 19S proteasome subunits is not limited to functions on chromatin. A recent study found 19S complex was abundant near brain synapses where it regulates synaptic proteins that control excitatory synaptic transmission, independent of the full 26S proteasome complex (96). Interestingly, we show proteasome inhibition, which predominantly increases the expression of 19S subunits, also results in open chromatin at regions that code for genes involved in neurogenesis. Altogether, 19S ATPase subunits enrichment at cell type specific enhancesomes, suggests subunit association with chromatin, which could be a potential mechanism for the increase DNA accessibility observed at distal cis-regulatory elements, when proteasome is inhibited.

Time constraints in this study do not allow us to pursue these questions, but future research may be necessary to address these potential limitations.

## DATA AVAILABILITY

The datasets supporting the results and conclusions of this article are available in the GEO repository, GEOXXXXX.

## SUPPLEMENTARY DATA

Supplementary data are available at NAR Cancer Online.

## AUTHOR CONTRIBUTIONS

Conceptualization, HKK and TKA; Methodology, HKK; Investigation, HKK; Formal Data Analysis, BDD, JMW, HKK; Visualization, HKK, BDD, JMW.

Writing – Original Draft, HKK; Writing – Reviewing & Editing, HKK, BDD, JMW and TKA. Reviewing & Editing Final; HKK, BDD, JMW and TKA; Funding Acquisition, TKA

## ACKNOWLEDGEMENTS

We would like to thank Drs. Sara Grimm, David Fargo, and Paul Wade at NIEHS for critical review of the manuscript. We thank Archer Lab members for discussions relating to the project and the NIEHS Epigenomics Core Laboratory for their next generation sequencing expertise.

---

**Figure S1.**
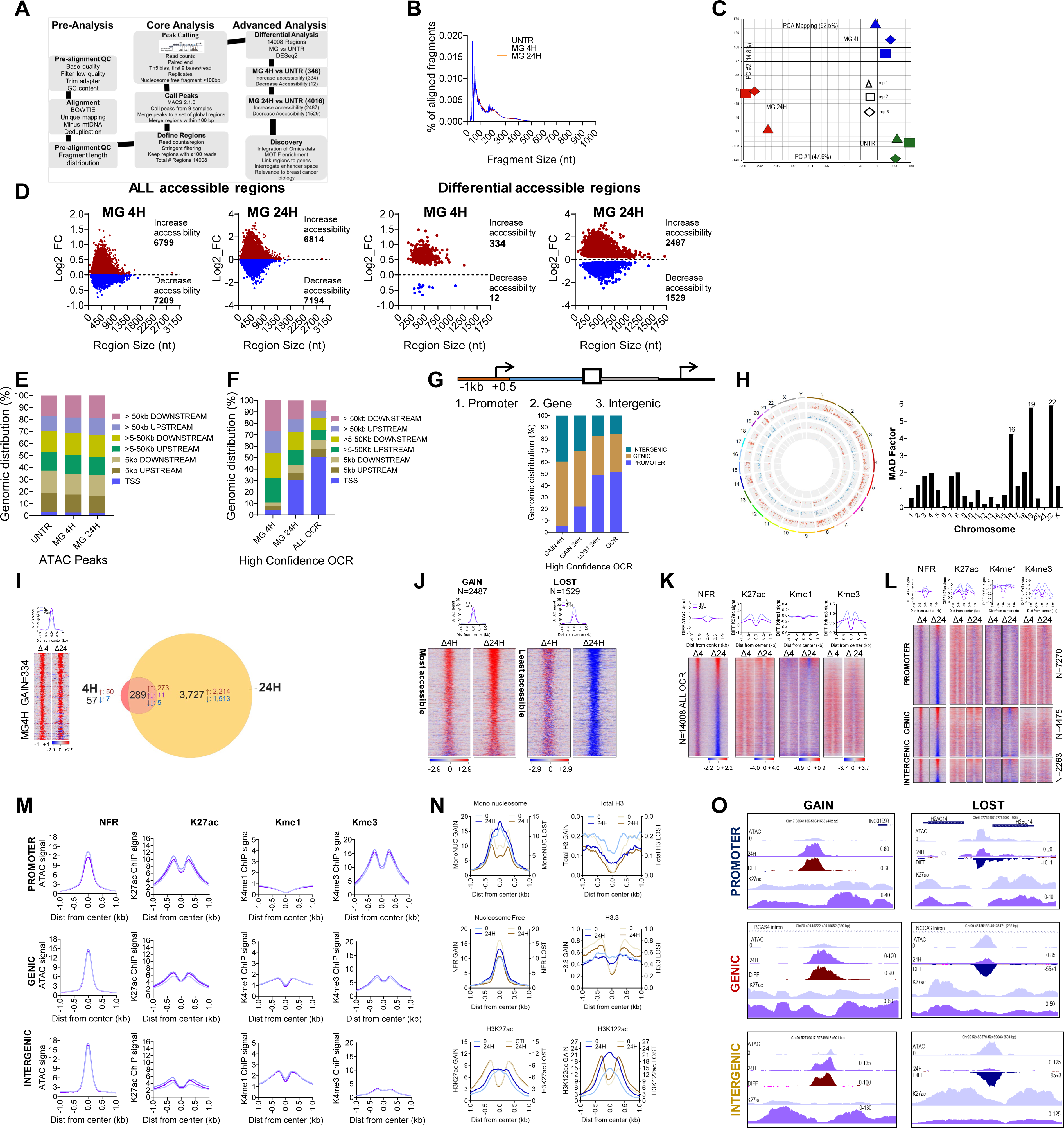
Systematic analysis of MG132 Differential Open Chromatin Regions (DOCRs) A) Flowchart describing the computational analysis pipeline used to define DOCRs from ATAC-seq. Major steps of analysis are highlighted. **B)** Graph showing insert size distribution of ATAC-seq fragments. **C)** Principal component analysis of ATAC-seq reads showing clear separation among treatment conditions, UNTR (Control, green), MG4H (blue), MG24H (red) are represented on two principal components with PC1 on x-axis and PC2 on y-axis. Three biological replicates per treatment condition are represented (rep1, rep2, rep3). PCA is based on ATAC signal, Tn5 transposase sensitive reads derived from 78, 380 high-confidence regions (FDR 0.05) with at least 10 reads in the 9 samples. **D)** Scatter plots showing the relationship between genomic size of accessible region (nt) and change in chromatin accessibility in MG132 treated samples (log2FC). ALL accessible regions (OCR) 4H (left) and 24H (right) and DOCRs that significantly increase or decrease chromatin accessibility at 4H (left) and 24H (right). **E)** Graph showing genomic distribution of accessible (ATAC) peaks in each treatment condition. **F)** Graph showing genomic distribution of MG132 high confidence open chromatin regions (OCRs) used in the analysis **G)** Graph showing distribution of high confidence OCRs by genomic category. TOP: Schematic defining genomic categories. **H)** Circos plot showing chromosome distribution of MG132 DOCRs. The outer ring represents reference chromosomes 1 through 22 in clockwise orientation. X axis denotes chromosome demarcation, y axis dots are DOCRs ranked by log2FC from 0 to max 2 change in accessibility. From outer circos ring Chromosome, 24H increase (blue), 24H decrease (red), 4H increase (blue), 4H decrease (red). **Right Graph:** Graph showing the Median Absolute Deviation (MAD Factor) indicating variability in DOCR chromosome distribution **I)** Chromatin accessibility of 334 GAIN DOCRs in cells treated for 4H. Heatmaps representing differential ATAC signal (NFR) and metaplot (Top) representing ATAC signal coverage at genomic regions that change accessibility at 4H, the lines represent signal in control, 0, 4H and 24H treated cells: **Right Panel:** Venn diagram showing the overlap of genomic regions that increase or decrease chromatin accessibility in cells treated for 4 and 24H. Numbers represent DOCRs in each class, arrows represent direction of change in accessibility (red, GAIN), blue, LOST). **J)** Heatmaps and metaplots (top) representing ATAC signal (NFR) at DOCRs that <u>GAIN</u> (2487, **Right Panel**) and <u>LOST</u> (1529, **Left Panel**). Heatmaps show differential accessibility (NFR), metaplots show ATAC signal in each treatment sample. **K)** Heatmaps and metaplots (TOP) showing differential NFR, H3K27ac, H3K4me1 and H3K4me3 signal at ALL OCR tested (14008) at 4 and 24H. **L)** Heatmaps and metaplots showing differential NFR, H3K27ac, H3K4me1 and H3K4me3 signal at ALL OCRs split in genomic categories, PROMOTER (7270), GENE (4475) and INTERGENIC (2263) regions. **M)** Metaplots showing chromatin state for all OCR PROMOTER (7270), GENE (4475) and INTERGENIC (2263). **N)** Metaplots showing distinct chromatin states of GAIN and LOST promoter DOCR. Unique chromatin architecture reflected by the unimodal and bimodal profiles of mono nucleosome signal, NFR, Total H3, H3.3, H3K27ac, H3K122ac. **O)** Browser tracks showing examples of GAIN and LOST DOCRs. Tracks show read coverage of chromatin accessibility (NFR), differential read coverage of accessibility (DIFF, GAIN=red, LOST=blue) and H3K27ac Track colors: Control (0), MG24H (24H).

**Figure S2.**
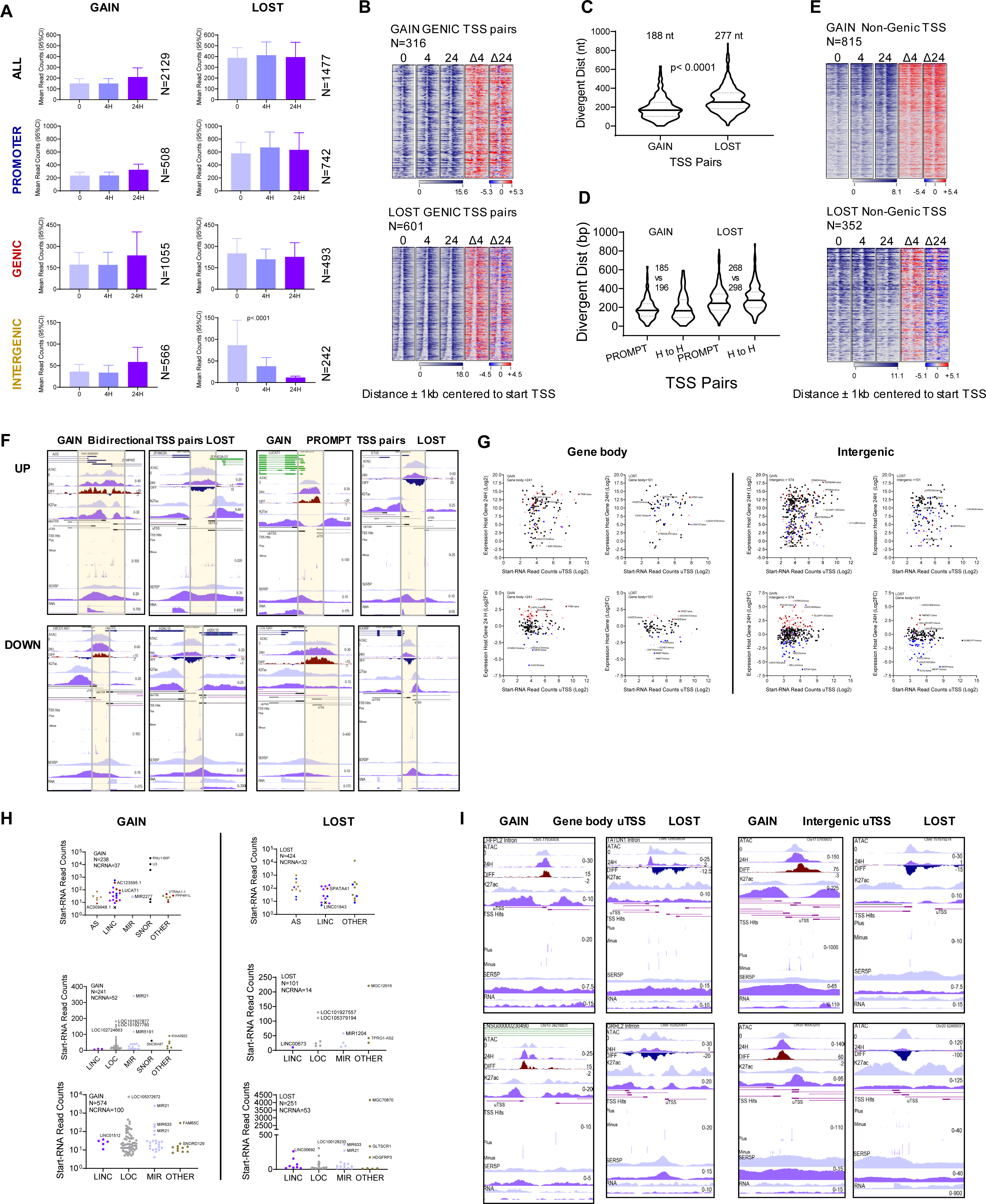
DOCRs are sites of RNAPII transcription. **A)** Graph representing RNA expression (RNA-seq) read counts overlapping DOCRs that significantly <u>increase (GAIN)</u> and <u>decrease (LOST)</u> accessibility. Reads from RNA-seq are within the region +/- 250 bp of the DOCR center. **B)** Heatmap showing average H3K27ac signal near genic TSS pairs within GAIN and LOST regions. Signal is ranked based on distance between TSS pairs. N is the number of genic TSS within each DOCR **C)** Violin plots comparing the distance between divergent GAIN and LOST TSS pairs. **D)** Violin plots comparing the distance between bidirectional (head-to-head) vs PROMPT TSS pairs. **E)** Heatmap showing H3K27ac signal at nongenic TSS. H3K27ac signal is sorted descending order based on total start-TSS read signal. N is the number of nongenic TSS within each DOCR. **F)** Browser tracks showing examples of bidirectional (head-head) and PROMPT TSS pairs around GAIN and LOST DOCRs. Tracks show read coverage of chromatin accessibility (NFR), differential NFR (DIFF), H3K27ac, start RNA hits, strand, Ser5P, RNA expression. Chromosome start coordinate for each TSS is shown. Names of TSS gene pairs are indicated. Track Control (0), MG24H (24H). Box highlights DOCR region and TSS hits. **G)** Scatter plot showing correlation between expression of nongenic TSS (start-seq) and closest gene (RNA-seq) read counts. **H)** Graph showing expression of different classes of noncoding RNAs nearby genic and nongenic TSS. **I)** Browser tracks showing examples of nongenic TSS in the gene body and intergenic GAIN and LOST DOCRs. Tracks show read coverage of chromatin accessibility (NFR), Differential NFR (DIFF), H3K27ac, start RNA hits, strand, Ser5P and RNA expression. Chromosome start coordinate for each TSS is shown. Gene names closest to the nongenic TSS in gene body are indicated. Tracks: Control (0), MG24H (24H).

**Figure S3.**
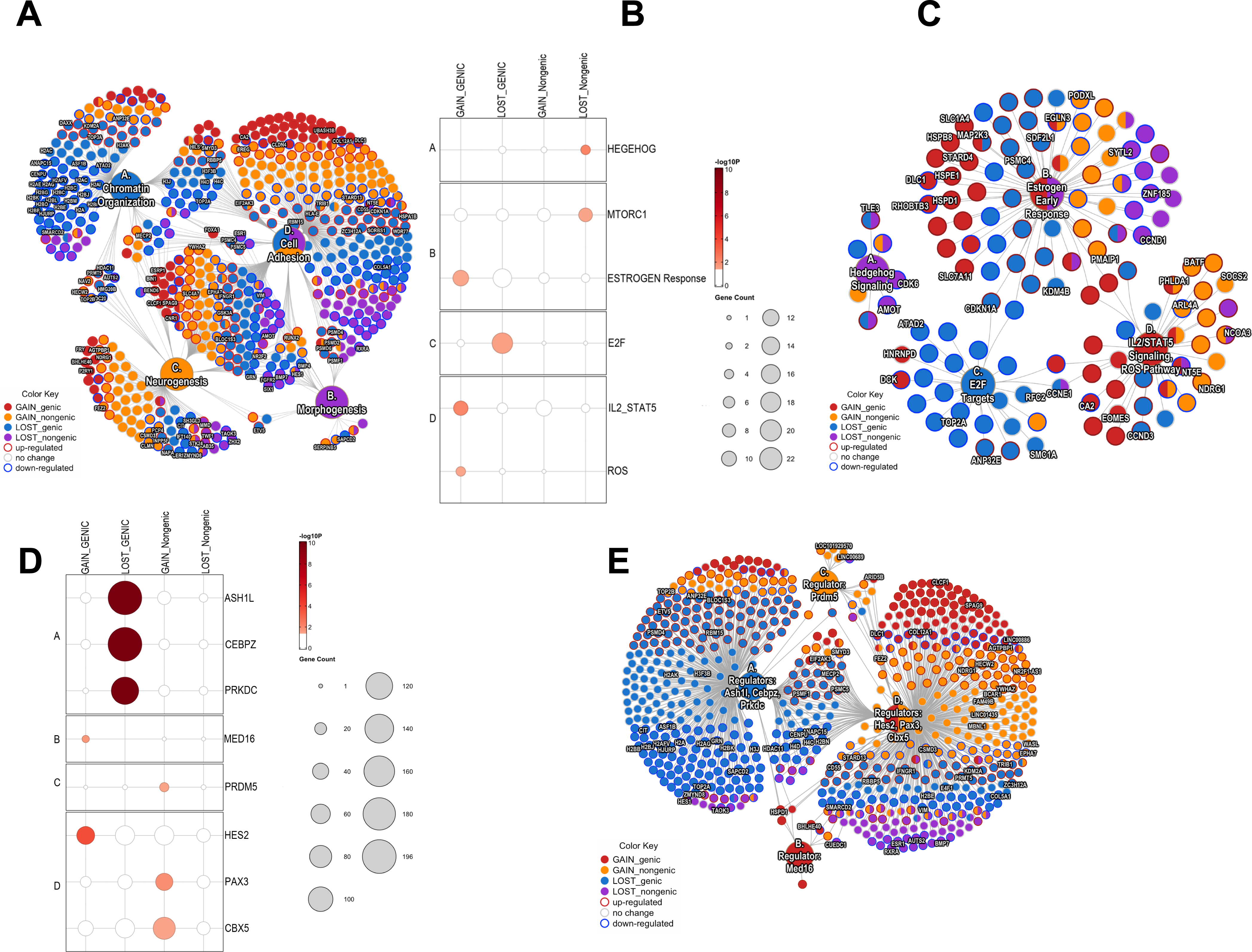
related to. **Figure 3. Biological processes affected by MG132 treatment A)** cNet-plot showing GO biological processes that were significantly enriched for genes closest to genic and nongenic TSS. The color of gene node represents the most enriched DOCR as per legend, genes in each node are organized based on direction of expression, border color of the dot identifies genes UP (red) or DOWN (blue) (log2FC ≤ 1, FDR=0.05) in cells treated for 24H. **B)** Enriched hallmark pathways. Dot plot showing the degree of enrichment for genes closest to DOCRs split by category. Dot size represents the number of genes in each hallmark pathway. **C)** cNet-plot showing hallmark pathways enriched for genes closest to genic and non-genic TSS. Key genes in each pathway are shown **D)** Transcriptional cofactors predicted to regulate genes closest to genic and nongenic TSS. Dot plot showing the degree of enrichment for genes closest to DOCRs split by category. Dot size represents the number of genes predicted to be regulated by each cofactor as determined from GTRD (gene transcription regulation database. **E)** cNet-plot showing GTRD factors whose targets are enriched for genes closest to genic and non-genic TSS.

**Figure S4.**
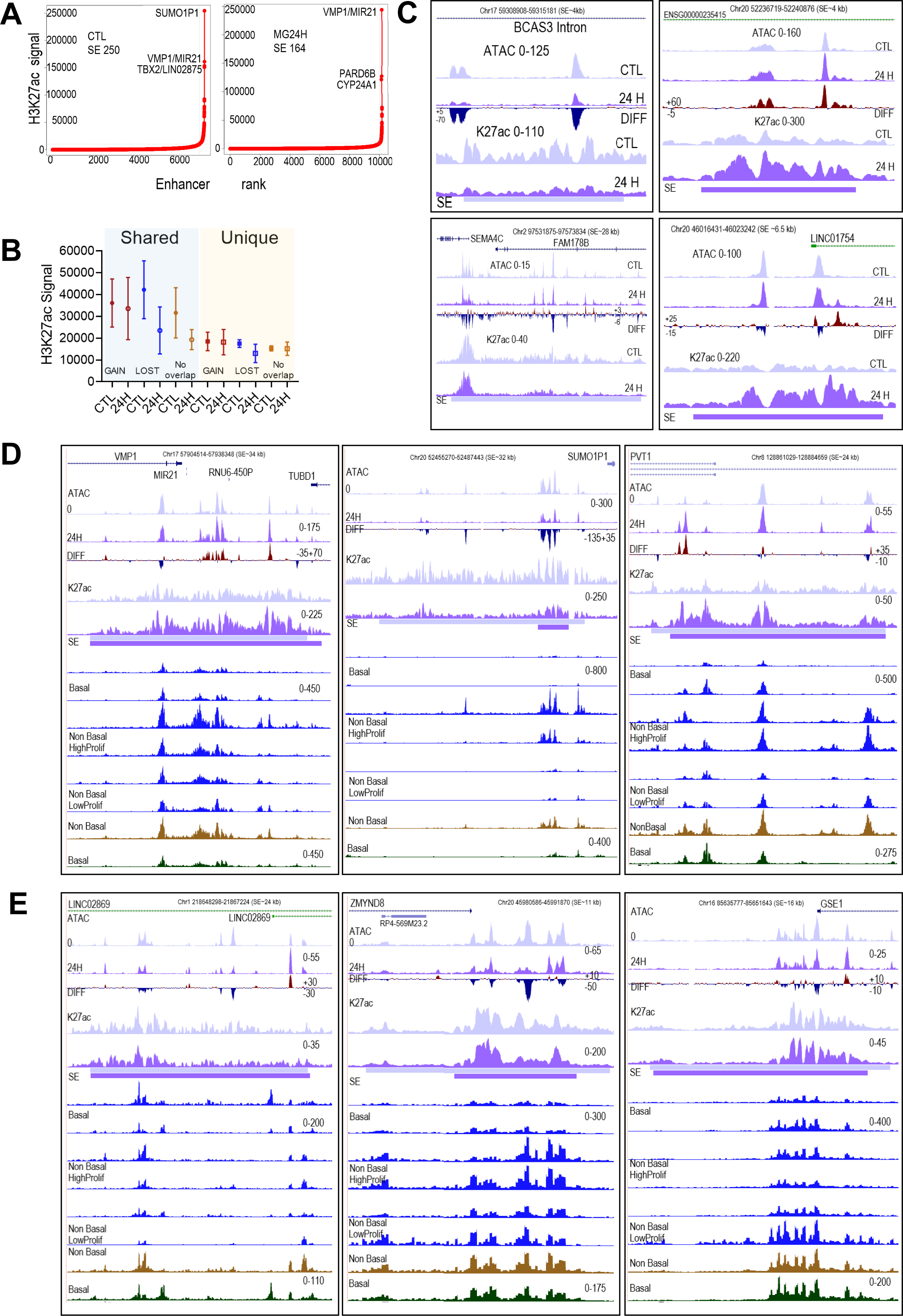
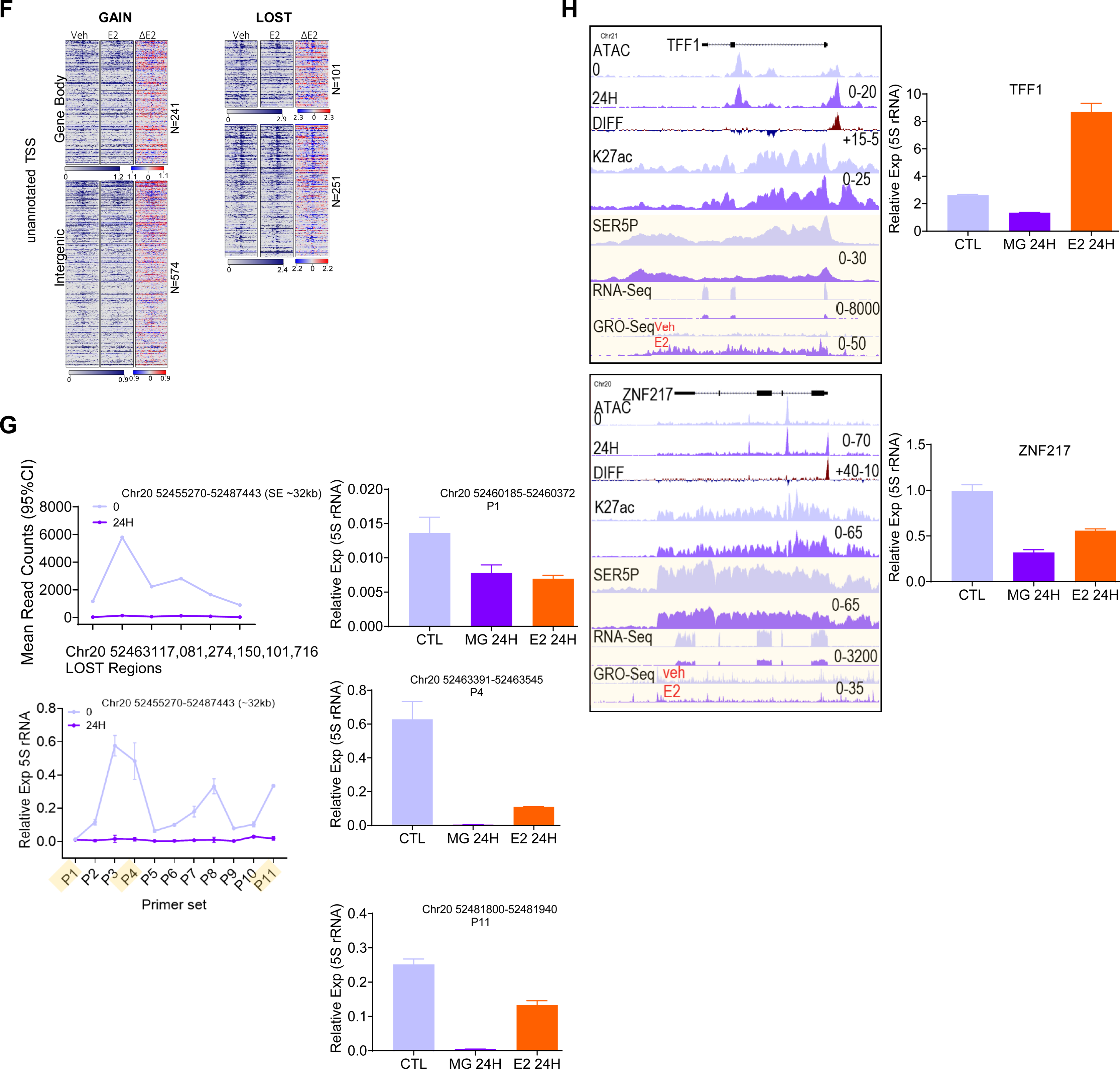
related to. Figure 4**. MG132 DOCRs overlap SE biologically relevant in breast cancer tumors. A)** Identification of super enhancers from control and MG132 treated MCF-7 cells. Enhancer regions are plotted in an increasing order based on normalized H3K27ac signal. Super-enhancers are defined as the population of enhancers above the inflection point of the curve. Examples of closest TSS to 3 top SEs in each class are shown **B)** Graph showing the level of H3K27ac signal observed at shared and unique super enhancers that overlap or do not overlap with DOCRs. **C)** Browser tracks showing examples of SEs 1) BCAS3 intron unique to control and overlaps with LOST DOCR; 2) Chr20 desert unique to MG24H, overlaps with GAIN; 3) SEMA4C/FAM178B unique to control and LINCO1754 unique to MG24H do not overlap with DOCRs. Tracks show read coverage of chromatin accessibility (NFR), differential NFR (DIFF), H3K27ac, and SE region. SE chromosome start coordinates are shown on top. Tracks: Control (0), MG24H (24H). **D)** SE show individual tumor heterogeneity in DNA accessibility. Browser tracks of VMP1/MIR21 (Chr 17), SUMO1P1 (Chr 20) and PVT1 (Chr 8), SE regions are representative examples, cluster A, B and D. Tracks show read coverage of chromatin accessibility (NFR), differential NFR (DIFF), H3K27ac, SE, normalized ATAC signal for individual tumors showing High or Low proliferation, average normalized ATAC signal for non-basal (brown) and basal (green) breast tumors. Tracks: Control (0), MG24H (24H). **E)** Browser tracks of C1orf143 (LINC02869, Chr1), ZMYND8 (Chr 20), GSE1 (Chr 16), SE regions are representative examples, cluster C, E and F. **F)** Nongenic DOCR transcription is responsive to E2 treatment. Heatmaps showing GRO-seq signal at nongenic TSSs that overlap GAIN (left) and LOST DOCRs (right) in cells treated with vehicle (veh) and estradiol (E2). **G)** MG132 and E2 repress transcription of SUMO1P1 SE region. Graph showing RNA-seq expression (read counts) of the DOCR regions that overlap with the SUMO1P1 SE. **Bottom**: Graph showing RNA expression from the SE region, x-axis shows the primer sets used qPCR analysis. Graph showing RNA expression of selected regions within SE in cells treated with MG132 and E2. Primer genomic coordinates are indicated **H)** Tracks showing chromatin state and transcription features of TFF1 and ZNF217, representative genes induced and repressed by E2. Graphs show RNA expression TFF1 and ZNF217. RNA expression normalized to 5S rRNA.

